# A doublecortin-domain protein of *Toxoplasma* and its orthologues bind to and modify the structure and organization of tubulin polymers

**DOI:** 10.1101/623140

**Authors:** Jacqueline M. Leung, Eiji Nagayasu, Yu-Chen Hwang, Jun Liu, Phillip G. Pierce, Isabelle Q. Phan, Robin A. Prentice, John M. Murray, Ke Hu

**Affiliations:** Department of Biology, Indiana University, Bloomington, IN, 47405, USA; Department of Infectious Diseases, Division of Parasitology, Faculty of Medicine, University of Miyazaki, Miyazaki, Japan; Nikon Instruments Inc., Melville, New York, 11747, USA; UCB, and Seattle Structural Genomics Center for Infectious Disease, Bainbridge Island, WA, 98110; Center for Global Infectious Disease Research, Seattle Children’s Research Institute, and Seattle Structural Genomics Center for Infectious Disease, Seattle, WA, 98109, U.S.A; University of Washington, Brotman Bady Institute, Seattle, WA 98195, U.S.A

## Abstract

TgDCX is a doublecortin-domain protein associated with the conoid fibers, a set of strongly curved non-tubular tubulin-polymers in *Toxoplasma*. TgDCX deletion impairs conoid structure and parasite invasion. TgDCX contains two tubulin-binding domains: a partial P25-alpha and the DCX/doublecortin domain. Orthologues are found in apicomplexans and their free-living relatives *Chromera* and *Vitrella*. We report that isolated TgDCX-containing conoid fibers retain their pronounced curvature, but loss of TgDCX destabilizes the fibers. We crystallized and determined the 3D-structure of the DCX-domain, which is similar to those of human doublecortin and well-conserved among TgDCX orthologues. However, the orthologues vary widely in targeting to the conoid in *Toxoplasma* and in modulating microtubule organization in *Xenopus* cells. Several orthologues bind to microtubules in *Xenopus* cells, but only TgDCX generates short, strongly curved microtubule arcs. EM analysis shows microtubules decorated with TgDCX bundled into rafts, often bordered on one edge by a “C”-shaped incomplete tube. A *Chromera* orthologue closely mimics TgDCX targeting in *Toxoplasma* and binds to microtubules in *Xenopus* cells, but does not generate arcs or “C”-shaped tubes, and fails to rescue the defects of the TgDCX-knockout parasite. These observations suggest that species-specific features of TgDCX enable it to generate strongly curved tubulin-polymers to support efficient host-cell invasion.

## INTRODUCTION

Microtubules are ubiquitous among eukaryotic cells. These polymers are formed of tubulin subunits, which are extraordinarily well-conserved. The vast majority of microtubules in the eukaryotic systems examined to date are formed of 13 protofilaments. However, there are a number of exceptions. Microtubules with atypical numbers of protofilaments have been found in several organisms [for reviews, see (Chaaban and Brouhard, 2017; Davis and Gull, 1983; Dustin, 1984)]. The B-tube in the doublet microtubule in the flagellum is an incomplete tube coupled with the 13-protofilament A-tube (Haimo and Rosenbaum, 1981). In the unicellular human parasite *Toxoplasma gondii*, a tubulin polymer that deviates significantly from the tube-form is found in the conoid (Hu et al., 2002). The conoid is a cytoskeletal complex of 14 fibers formed into a truncated cone. Each conoid fiber is a strongly curved [Figure 1A, (Hu et al., 2002)] folded ribbon formed of 8-9 tubulin protofilaments and follows a trajectory of a left-handed spiral. The unusual structure of the conoid fibers is not dictated by tubulin itself, because the tubulin subunits in these fibers are identical in amino acid sequence to those forming canonical microtubules elsewhere in the parasite (*e.g.* the cortical microtubules, intra-conoid microtubules, and microtubules in the mitotic spindle). The conoid is a motile organelle. As part of the *T. gondii* lytic cycle, which is the fundamental pathogenetic event in toxoplasmosis, in response to the signal of elevated intracellular [Ca^2+^], the parasite activates motility to exit its lysed host cell and extends the conoid. Extension is accompanied by a change in the shape of the conoid as it becomes less conical and more cylindrical, while the pitch of the conoid fibers becomes steeper (Hu et al., 2002) (Figure 1A).

**Figure 1.**
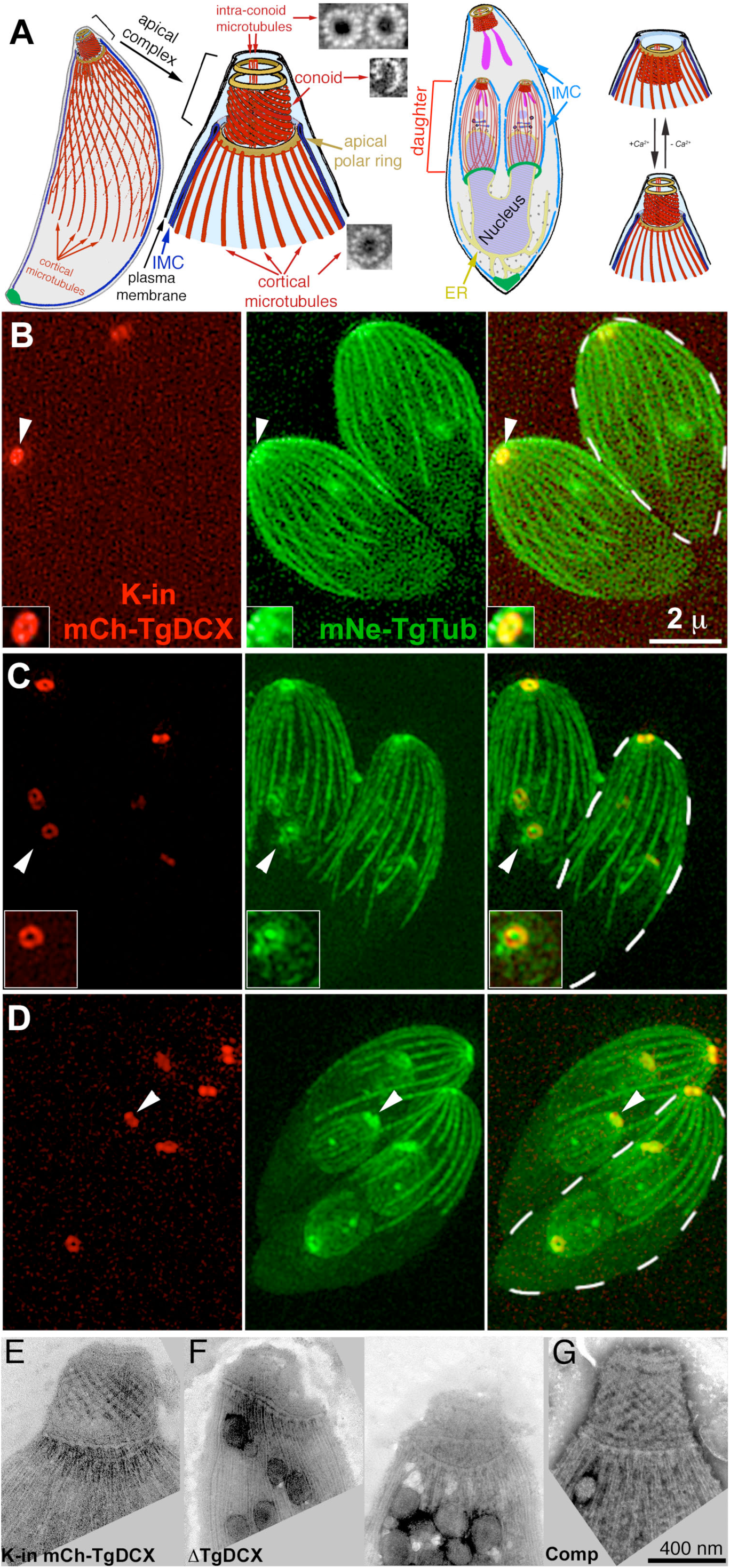
Conoid architecture and endogenous TgDCX localization (A) Diagrams of the *T. gondii* cytoskeleton [modified from (Nagayasu et al., 2016)], in which several tubulin containing structures (22 cortical microtubules, 2 intra-conoid microtubules, and 14 conoid fibers) are highlighted in red. EM images of a cross-section of each of those polymers (Hu et al., 2002) are also shown. Shown in brown are several rings mentioned later in the text. The apical polar ring is the origin of the 22 cortical microtubules. A complex structure (the preconoidal rings, unlabeled), rich in intricate detail, lying at the apical end of the conoid, is portrayed in this cartoon as two featureless rings. IMC: Inner Membrane Complex. A replicating parasite is also shown, with daughter parasites being built inside the mother. The cortical microtubules of the adult are omitted for clarity. On the right, a cartoon shows how the conoid responds to increasing [Ca^2+^] by extending and changing its shape. (B-D) Z-projections of SIM images of mCherryFP-TgDCX (*red*, “K-in mCh-TgDCX”) knock-in parasites (Nagayasu et al., 2016) expressing mNeonGreenFP-β1-tubulin (*green*, mNe-TgTub). (B) Two interphase adult parasites. One adult is outlined with a dashed white border. The arrowhead indicates the apical complex of one parasite, shown 2x enlarged and contrast enhanced in the inset. Tubulin and TgDCX are co-localized in the conoid, appearing as an annulus with a ∼ 0.2 µm central opening. (C) Two dividing parasites at an early stage of daughter formation, with two daughters in each adult. One of the developing daughter’s apical complex is indicated by the arrowhead, and enlarged 1.5x in the inset. (D) Parasites at a later stage of daughter formation. The daughter apical complexes (*white arrowhead*) are nearly mature, and daughter cortical microtubules have grown to ∼ 1/3 of their length in the adult. (E-G) Electron microscope (EM) images of the conoid region of negatively stained whole-mount *mCherryFP-TgDCX* knock-in (E, “K-in mCh-TgDCX”), *TgDCX* knockout (F, “ΔTgDCX”) parasites (two images), and a complemented line generated by transfecting the *TgDCX* knockout parasite with a plasmid driving expression of TgDCX-EGFP (G, “Comp”). The conoids are shorter, distorted, and disordered in the *TgDCX* knockout parasites (F) compared to their parental strain in (E), but supplying TgDCX completely restores conoid structure (G).

To search for the non-tubulin components responsible for the formation and maintenance of the unusual structure of the conoid fibers, a conoid-enriched fraction of the *Toxoplasma* cytoskeleton was analyzed by mass spectrometry, which revealed many novel candidate components (Hu et al., 2006). One of the proteins identified was TgDCX, localized exclusively along the conoid fibers (Nagayasu et al., 2016). TgDCX contains a doublecortin (DCX) domain and a partial P25α domain, two predicted tubulin-binding domains normally found separately in the proteins of mammalian neuronal cells. Human doublecortin, which contains two DCX domains, cooperatively binds to the groove between protofilaments (Bechstedt and Brouhard, 2012; Fourniol et al., 2010; Moores et al., 2004). P25α (also known as tubulin polymerization perturbing protein, TPPP) also binds to tubulin and modifies the structure of microtubules (Hlavanda et al., 2002). Mutations in the human *doublecortin* gene lead to gross disturbances in the architecture of the cerebral cortex: the double cortex syndrome in females and lissencephaly in males, due to perturbed neuronal migration during development (Caspi et al., 2000; Gleeson et al., 1998; Kato et al., 1999). The partial P25α domain of TgDCX has in fact significant homology to a consensus DCX domain (30% similarity 22% identity), so it was reasonable to hypothesize that the P25α and DCX domains of TgDCX function in a similar way as the two DCX domains of doublecortin. The linker region between the partial P25α domain and the DCX domain of TgDCX (44 aa residues) is slightly longer than that between the two DCX domains of doublecortin (40 residues).

Human doublecortin has been proposed to stabilize the 13-protofilament microtubule structure in preference to other architectures that readily form when tubulin is polymerized in the absence of other proteins (Moores et al., 2004). This may be due to its ability to sense curvature in the protofilaments. It is intriguing that it seems to prefer strongly curved regions of microtubules in the presence of paclitaxel, and straight microtubules in the absence of paclitaxel (Bechstedt and Brouhard, 2012; Ettinger et al., 2016). Given this dramatic curvature-sensitive interaction of DCX domains with tubulin, TgDCX is a prime candidate for the role of shaping conventional tubulin into unconventional highly-curved conoid fibers.

Although both the P25α and DCX domains are ubiquitous among metazoa, the presence of both domains in the same molecule appears to be almost exclusively an apicomplexan feature (Orosz, 2009; Orosz, 2016). This combination is found in all sequenced genomes of apicomplexans (*e.g. Toxoplasma gondii, Plasmodium* spp. and *Cryptosporidium* spp.), as well as their photosynthetic (*Chromera velia* and *Vitrella brassicaformis*) and parasitic (*e.g. Perkinsus* spp.) relatives in the superphylum Alveolata. Beyond Apicomplexans and their relatives, the P25α+DCX arrangement is found in a gene in the sequenced genome of *Trichoplax adhaerens* from the phylum *Placozoa*, a primitive metazoan (Srivastava et al., 2008). Genomic sequence data from Eumetazoa occasionally appear with purported “apicortins”, but these likely result from contamination with the DNA of an apicomplexan parasite infecting the Eumetazoan host (Orosz, 2018). An “apicortin” was also reported in the genome of the sperm whale (*Physeter macrocephalus)*, but this was likely due to infection of the whale by *Sarcocystis*, a common apicomplexan parasite of marine mammals. Given these examples, it cannot be excluded that the TgDCX orthologue reported in the *Trichoplax adhaerens* genome is from a marine microbe distantly related to the apicomplexans.

Previously we reported that TgDCX is specifically localized to the conoid in *Toxoplasma*, and that without TgDCX, the conoid structure becomes short and disordered, invasion of host cells is severely compromised, and the parasite forms many fewer and much smaller plaques relative to wild-type *Toxoplasma* (Nagayasu et al., 2016). We report here a more in-depth analysis of the conoid fiber architecture, and a comparison of TgDCX and its eight orthologues from *Plasmodium falciparum* (an obligate intracellular parasite)*, C. velia* and *V. brassicaformis* (free-living phototrophs), and *T. adhaerens* (an early diverging metazoan), utilizing localization *in vivo*, structural modeling based on the crystal structure of the DCX domain of TgDCX, and cross-complementation analyses in *Toxoplasma*. These comparisons revealed that neither microtubule binding nor conoid-targeting is sufficient for functional complementation. TgDCX alone has the unique ability of generating short, curved, and stable tubulin-containing polymers with a consistent curvature. It also recruits another conoid component to the microtubules in *Xenopus* cells. This suggests that the divergent regions of TgDCX are important for modulating its function, and likely related to its ability to generate and stabilize tightly curved tubulin polymers.

## RESULTS

### TgDCX likely stabilizes the conoid fibers

Proteins containing either the P25α or DCX domain are typically involved in interactions with microtubules (Bechstedt and Brouhard, 2012; Fourniol et al., 2010; Gleeson et al., 1999; Hlavanda et al., 2002; Moores et al., 2004). However, in *Toxoplasma*, TgDCX is specifically localized to the conoid fibers and is not associated with the canonical 13 protofilament (pf) microtubules that are present elsewhere in the same cell (Nagayasu et al., 2016). Figure 1B-D shows the localization of TgDCX in parasites whose single endogenous genomic copy of *TgDCX* has been replaced with *mCherryFP-TgDCX* (*i.e*., “mCherryFP-TgDCX knock-in” parasites) (Nagayasu et al., 2016). Colocalization with mNeonGreenFP-β1-tubulin shows that, in both interphase and dividing parasites (in which the daughters form inside the mother), mCherryFP-TgDCX is specifically concentrated in the conoids, but absent from all other tubulin-containing structures, such as the cortical microtubules or the centrosome. This localization is consistent with the conoid-specific structural defect (Figure 1E-G) when the *TgDCX* gene is deleted (Nagayasu et al., 2016).

To further characterize its structural role, we attempted to separate the TgDCX-containing conoid fibers from the rest of the cytoskeleton in wild-type (RH*Δku80Δhx*), *mCherryFP-TgDCX* knock-in, and *TgDCX* knockout parasites (Figure 2A-D). Preparations from the wild-type and knock-in parasites contain intact conoids (Figure 2B) as well as numerous free conoid fibers that are no longer associated with other structures in the apical complex (Figure 2A and 2C). The conoid fibers are always curved into short arcs. In the intact conoid, the fibers follow inclined paths on the surface of a cone (Figure 2E). The intersection of a cone with an inclined plane is an egg-shaped curve; *i.e*., a closed curve with only one symmetry axis and continuously changing radius of curvature. However, for a cone with the dimensions of the retracted conoid and for a plane inclined at the angle of the conoid fibers, the intersection is nearly elliptical, with major and minor axes of the ellipse that differ by ∼10%. The conoid fiber path covers much less than half of the circumference of the conoid, so in practice its shape can be closely approximated by an arc of a circle. The free conoid fibers all have similar arc length (355 ± 3.8 nm; sem, n=100), span the same arc angle (81.5 ± 1.2^°^), and have the same radius of curvature (253 ± 3.7 nm), indicating that these fibers are stable and their pronounced curvature is an intrinsic feature of their structure. In contrast, the fragments of cortical microtubules seen in the same image are all remarkably straight (Figure 2A). Note that it is not possible to bend a microtubule into an arc as tight as that of the conoid fibers: when forcefully bent, normal microtubules snap before reaching curvatures this pronounced (Amos and Amos, 1991).

**Figure 2.**
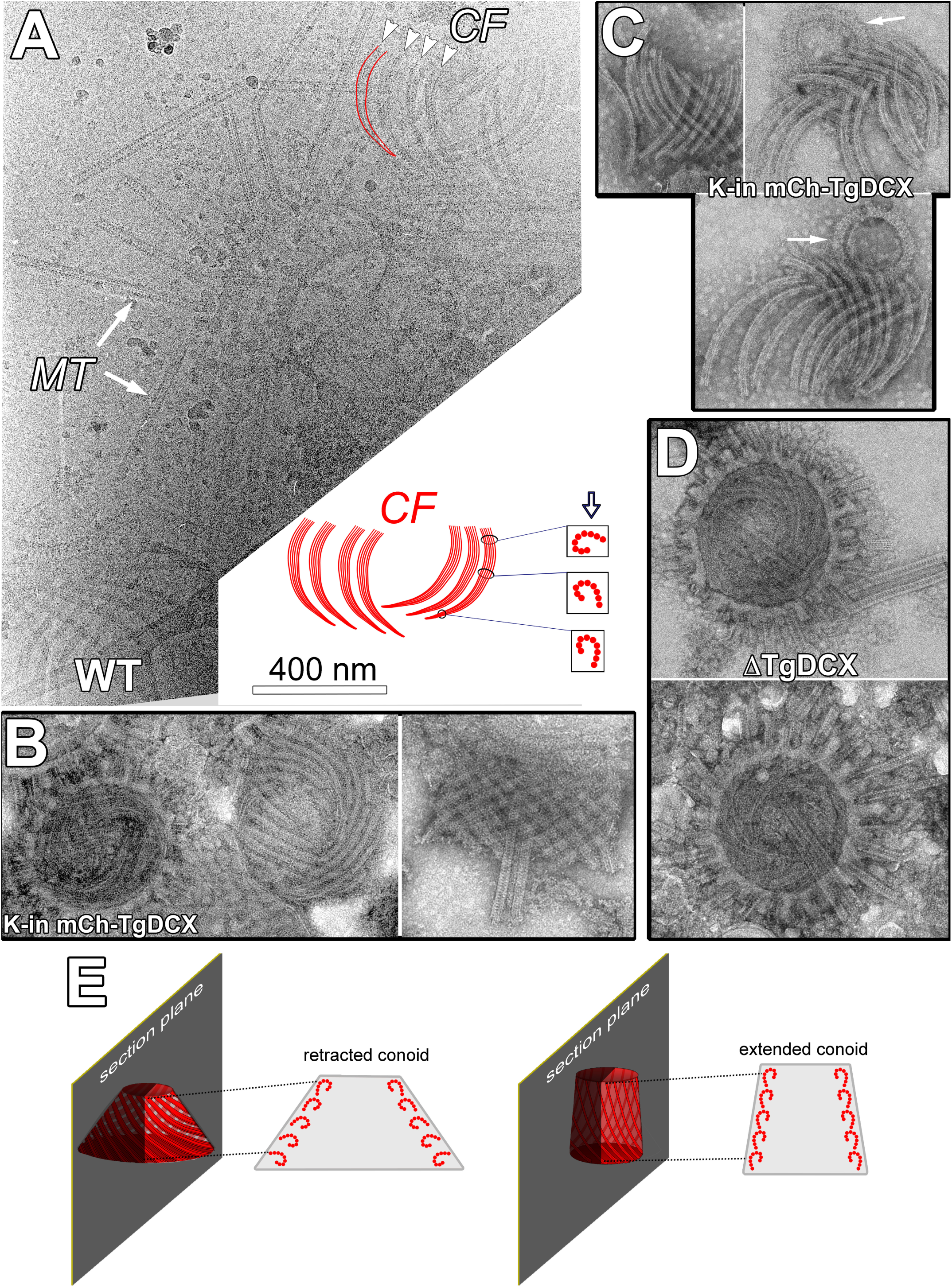
Conoid diagram and EM images of conoids isolated from wild-type, *mCherryFP-TgDCX* knock-in and *TgDCX* knockout parasites (A) CryoEM image of disassembled apical complexes from wild-type (“WT”) parasites. Several groups of conoid fibers (“*CF*”, *arrowheads*) and fragments of cortical microtubules (“*MT*”, arrows) are seen. Note that the cortical microtubules are straight, whereas the conoid fibers are uniformly curved. The conoid fibers appear to become wider along their length, and their protofilaments become clearer, indicating a twist in the fibers, as diagrammed in the cartoon. The cartoon represents the cluster of fibers in the upper right of the cryoEM image. The hollow arrow in the cartoon shows the direction of view in the EM image, and the boxes contain cross-sections of the fiber at the indicated locations. Near the apical end of the fibers (towards the bottom in the cartoon and the image), the direction of view yields a narrow fiber with protofilaments obscured by superposition. As the fiber twists along its length, its profile becomes wider, and there is decreasing superposition of protofilaments, giving the splayed appearance at the basal region of the fibers. (B) End on (left) and side views (right) of negatively stained isolated conoids from *mCherryFP-TgDCX* knock-in parasites (“K-in mCh-TgDCX”). In the left image, the apical polar ring with stumps of broken cortical microtubules encircles the conoid. A second conoid, almost completely disassembled, is also seen. In the right image, the two intra-conoid microtubules are seen projecting through the conoid, which is detached from the apical polar ring. (C) Three examples of disassembled conoids isolated from *mCherryFP-TgDCX* knock-in parasites (“K-in mCh-TgDCX”). All 14 of the fibers that formed each conoid are seen. Arrows: preconoidal rings, which often remain attached to the apical ends of the fibers. (D) End on views of conoids isolated from *TgDCX* knockout parasites (“ΔTgDCX”). The conoids are encircled by the apical polar ring with attached fragments of cortical microtubules. Isolated conoid fibers or conoid detached from the apical polar ring were never observed in preparations from the *TgDCX* knockout parasite. (E) Diagram illustrating the changing geometry of the TgDCX-containing fibers of the conoid. Extension of the conoid through the apical polar ring, which occurs as the parasites reactivate motility and exit their lysed host cell, is accompanied by a change in conoid shape from more conical to more cylindrical. The structural implications, for the conoid fibers, that follow from this change in overall shape are described in the Discussion. For clarity, the change in fiber orientation has been exaggerated in the diagram. Note also that the diagrams here are oversimplified for clarity: in an untilted sagittal section as diagrammed, the fiber profiles cannot be clearly seen. In order to make the profiles visible, the section must be tilted in the microscope by plus (to see the profiles on one side) or minus (to see the profiles on the other side) the pitch angle of the fibers. See Fig 6 in (Hu et al., 2002) for a demonstration.

Close examination of conoid fiber images such as those in Figure 2A commonly reveals increased visibility of the protofilaments and a significant broadening of the fibers at their basal ends (*i.e.* the end closest to the apical polar ring when the conoid is extended) compared to the apical ends (*i.e.* the end closest to the preconoidal rings; *cf.* Figure 1A). Using images of negatively stained conoid fibers, for which the contrast is higher than for cryoEM, we measured an average increase in apparent width of 5.4 nm in going from the apical to basal regions of the fibers (95% confidence interval 4.4 to 6.4 nm; significant with p<0.0001). As the fibers have an asymmetric cross-section, similar to a dot-less elongated question mark (“?”) or an inverted “J”, their apparent width depends on the direction of view. Thus, the broadening at the basal end of the fibers is probably indicative of an inherent twist in the fiber, effectively changing the direction of view as one proceeds from apical to basal end (see the diagram in Figure 2A). For reasons to be described later, we propose that this is a left-handed twist (*See Discussion*).

Disordered conoid fibers can be observed in some intact conoids isolated from *TgDCX* knockout parasites (Figure 2D). However, unlike preparations from wild-type or *mCherryFP-TgDCX* knock-in parasites, free conoid fibers are not seen. We used the apical polar ring as a reference for calibrating the frequency of isolated conoid fiber visualization. There should be one intact conoid or cluster of conoid fibers present for each apical polar ring, if both structures survived the preparation for EM imaging equally well. In images of negatively stained *mCherryFP-TgDCX* knock-in parasites, “empty” apical polar rings (i.e., lacking an associated conoid), and detached but nearly complete conoids, or conoid fibers in clusters, were commonly observed (Figure 2B-C). Systematically counting every recognizable structure on two typical grids gave 109 “empty” apical polar rings and 39 detached conoids/conoid fiber clusters. In contrast, images of DCX-knockout parasites treated in the same way contained no recognizable conoid fiber clusters nor intact conoids detached from the apical polar ring. Counting two grids yielded 101 “empty” apical polar rings, but zero detached conoids or conoid fibers, suggesting that the conoid fibers are less stable without TgDCX. Alternatively, it is also possible that the fibers are more difficult to separate from the rest of the conoid in the absence of TgDCX. However, the latter seems less likely, as one would have expected on occasion to encounter an isolated escaped fiber, or at least a fragmented conoid with fibers hanging loose, but these have never been seen.

### TgDCX promotes the generation and stabilization of curved microtubules in a heterologous system

To characterize the effect of TgDCX on the structure of tubulin polymers in the absence of any other *Toxoplasma* proteins, ideally one would begin with *in vitro* reconstitution experiments using purified tubulin and TgDCX. Unfortunately, this is not possible, as TgDCX protein *in vitro* is insoluble in buffers that are compatible with polymerization of tubulin (see *Methods*). As an alternate strategy, we characterized the interaction of these two proteins *in vivo* by expressing FP-tagged TgDCX in *Xenopus* S3 cells to determine the effect of TgDCX in the absence of any other *Toxoplasma* proteins. We found that TgDCX not only coats tubulin-containing polymers in this heterologous system but also drastically alters the organization of the microtubule cytoskeleton (Figure 3 and Supplementary movies1&2). In untransfected cells (*e.g.* in Figure 3B, Figure 4A, and Figure 5A), the microtubules are long and extended, and the radius of curvature for ∼ 55% of the population is greater than 20 µm (mean = 117 µm; sem 9.4 µm, n = 844, total length 7488 µm). In contrast, TgDCX expression generates many short and smoothly curved fibers. The mean radius of curvature of TgDCX associated microtubules is 4.7 µm (sem = 0.17 µm, n=599; total length 2621 µm) (Figure 3C). A small fraction of the TgDCX associated microtubules are “flexible” (*i.e.,* meander through the cytoplasm on idiosyncratic trajectories with mostly gentle curves, as do normal microtubules. Using a *Sindbis* virus that contains 240 copies of mCherryFP-tagged capsid protein as the calibration standard (Murray, 2017), the number of TgDCX molecules per micron of fiber length for this class of fiber is 1459 (sem = 131, n=70) (Figure 3D-E). In a canonical microtubule, there are 1625 dimers of tubulin per micron, so if this “flexible” class represents single microtubules, then 0.9 TgDCX molecules are bound per tubulin dimer. The number of DCX molecules per micron for the short, smoothly curved microtubule arcs is higher (mean = 4854) and more variable: the 5%-95% range is 1096 to 12,030 TgDCX molecules per micron, suggesting that the curved microtubules are bundles containing on average 3 microtubules, which is consistent with the EM observations to be described later. This type of TgDCX-coated fiber is resistant to depolymerization by nocodazole treatment (Figure 4).

**Figure 3.**
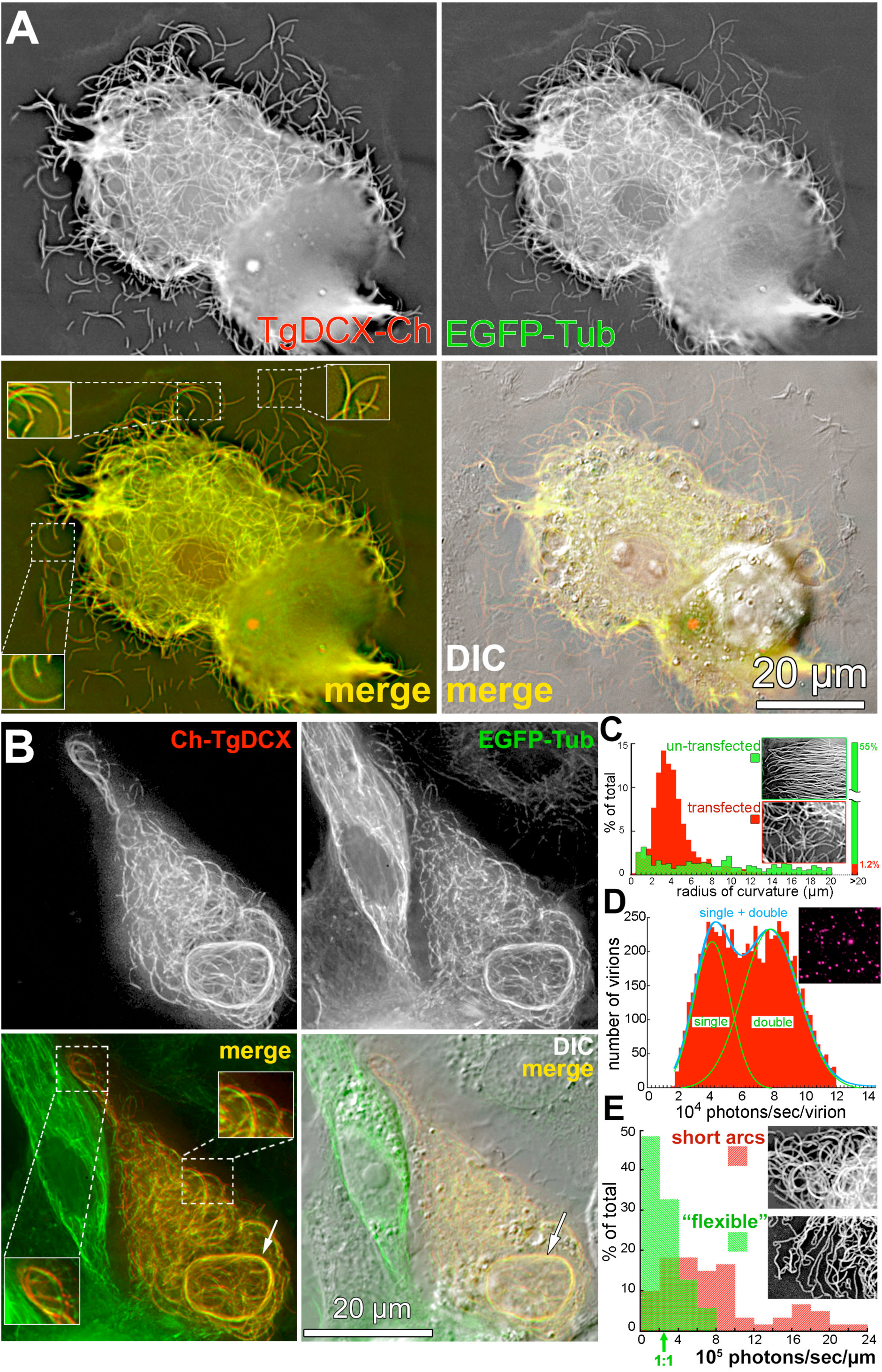
FP-tagged TgDCX generates curved microtubules in a heterologous system, *Xenopus laevis* S3 cells. (A) Deconvolved wide-field images of *Xenopus* S3 cells expressing EGFP-α-tubulin (*green*) and TgDCX-mCherryFP (*red*), showing that TgDCX-mCherryFP is localized to EGFP-α-tubulin containing fibers. TgDCX-mCherryFP expression drives the formation of curved fibers, many of which are short and share similar curvatures (also see Supplementary movie 1). The areas within the dashed outlines are enlarged 1.5x in the small white boxes. (B) Deconvolved wide-field images of *Xenopus* S3 cells expressing EGFP-α-tubulin (*green*) and mCherryFP-TgDCX (*red*), showing that, similar to TgDCX-mCherryFP, mCherryFP-TgDCX expression drives the formation of curved fibers (also see Supplementary movie 2). Arrows indicate microtubule bundles coated with mCherryFP-TgDCX. Note the fibers containing EGFP-tubulin only in the neighboring untransfected cell, which follow the meandering, gently curved, paths of typical cytoplasmic microtubules. Insets: 1.5x (C) Histogram of radii of curvature of microtubules in *Xenopus S3* cells, untransfected (green), or transfected with mCherryFP tagged TgDCX (red). Data include 844 microtubule segments (total length 7488 µm) for the untransfected cells and 599 segments (total length 2621 µm) for the transfected cells. All measurable microtubule segments in 8 representative untransfected cells and 7 transfected cells were included in the two histograms. Insets: fluorescent images of subregions of an untransfected (top) and a transfected cell (bottom). (D) Distribution of photon counts in images of *Sindbis* virus containing a capsid protein tagged with mCherryFP to be used as the fluorescence calibration standard for molecule counting. Y-axis: number of virions. X-axis: 10^4^ photons/sec/virion. The images commonly include both single virus particles and a variable proportion of unresolved pairs, and occasionally a small number of larger aggregates. The single and double particles can be distinguished as two peaks in the histogram of measured intensities. The smooth blue and green curves overlying the histogram show the least squares fit to a Gaussian distribution with two components. The empirical means (sem) for the two components derived from this fitting are 38318 (494) and 77406 (683) photons/sec/virion. Inset: fluorescent image of a field of mCherryFP tagged *Sindbis* virus particles. (E) Photon count distribution in images of mCherryFP tagged TgDCX associated with “flexible” microtubules (green) or short curved arcs (red). Arrow indicates the expected photon count for FP tagged TgDCX molecules to decorate a single microtubule in a 1:1 TgDCX to tubulin dimer ratio. Y-axis: % of total. X-axis: 10^5^ photons/sec/μm. Insets: fluorescent images containing “flexible” microtubules (bottom) or short curved arcs (top).

**Figure 4.**
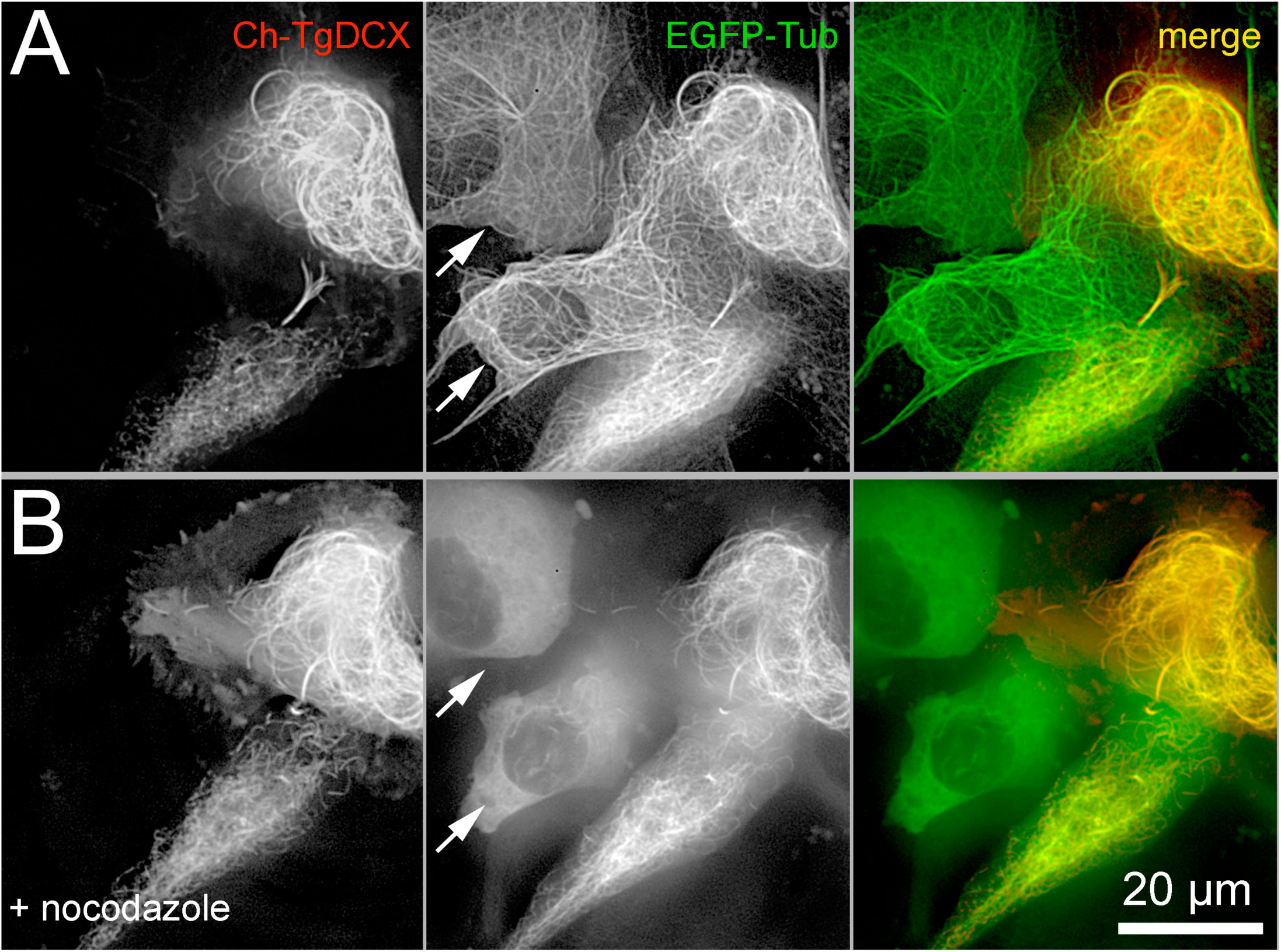
Curved fibers coated with mCherryFP-TgDCX are stable Deconvolved wide-field images of *Xenopus* S3 cells expressing EGFP-α-tubulin (*green*) and mCherryFP-TgDCX (*red*) before (A) and after (B) treatment with 16 µM nocodazole for 65 min. Note that the curved fibers coated with mCherryFP-TgDCX are resistant to depolymerization by nocodazole. Arrows indicate untransfected cells, expressing EGFP-α-tubulin only, in which virtually all microtubules have depolymerized.

**Figure 5.**
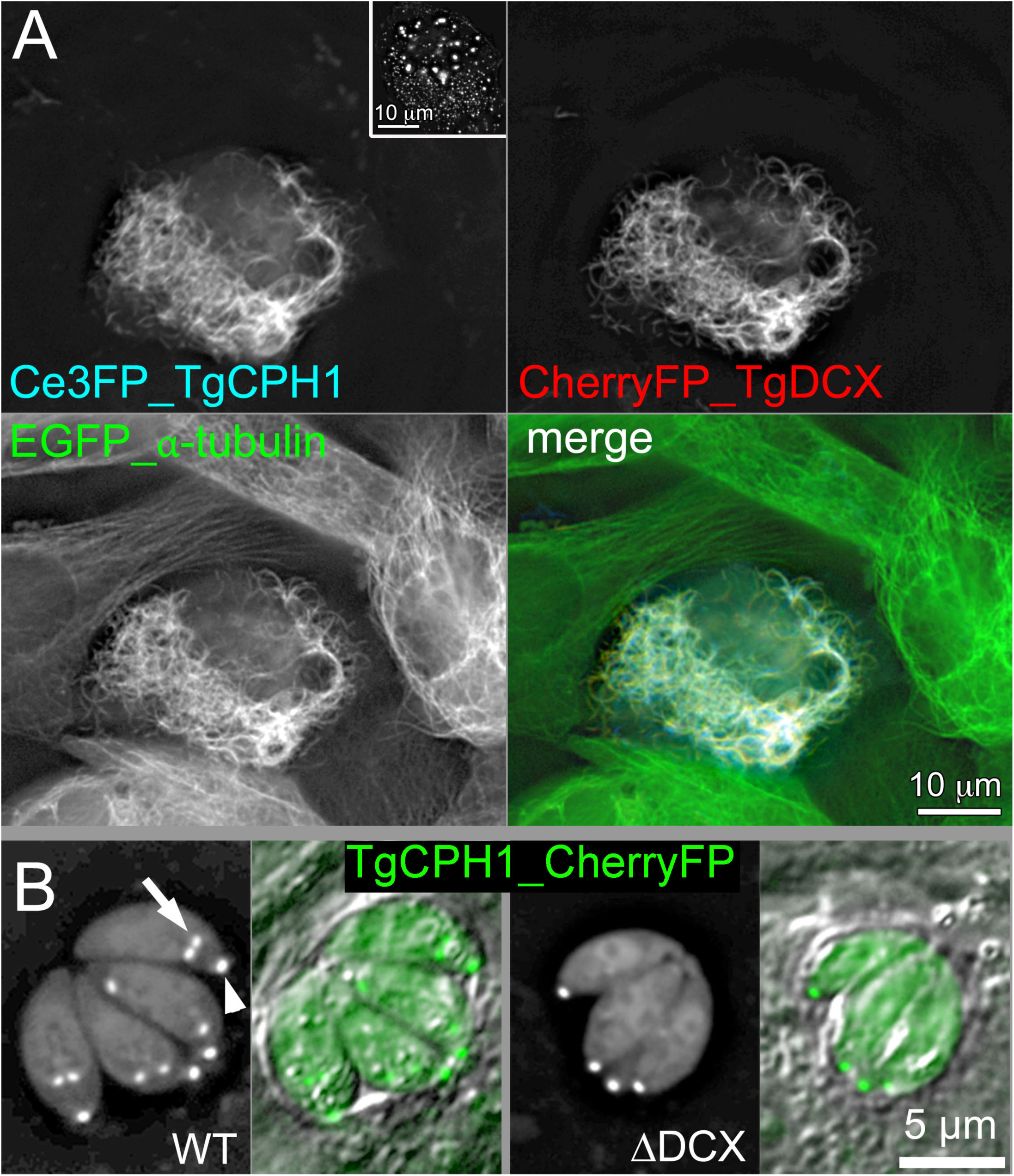
TgDCX bridges TgCPH1 to tubulin (A) Deconvolved wide-field images of *Xenopus* S3 cells expressing mCerulean3FP-TgCPH1, mCherryFP-TgDCX and EGFP-α-tubulin. Inset: a control *Xenopus* S3 cell that expresses mCerulean3FP-TgCPH1 but not TgDCX (also see Supplementary Figure 1). CPH1 on its own does not bind to microtubules in *Xenopus* cells, but is recruited to the microtubules by TgDCX. (B) RH*Δku80Δhx* (“WT”) and TgDCX knockout (ΔDCX) parasites expressing mCherryFP tagged TgCPH1. CPH1 localizes to daughters (arrows) and adult conoids (arrowheads) regardless of the presence of TgDCX.

The TgDCX induced arcs in *Xenopus* cells are much less sharply curved (mean radius of curvature 4.7 µm) than the conoid fibers in *Toxpolasma* (∼ 0.25 µm). TgCPH1, first identified as a candidate conoid protein in a previous proteomic screen (Hu et al., 2006), has been recently confirmed to be an important conoid component (Long et al., 2017). We therefore examined whether TgDCX and TgCPH1 act in concert. When expressed alone in *Xenopus* cells, TgCPH1 does not associate with microtubules, but is found in the cytosol, the nucleus, as well as in numerous vesicles (Supplementary Figure 1, Figure 5 inset). When co-expressed with TgDCX, however, TgCPH1 is recruited to the microtubules, slightly decreasing the radius of curvature of the arcs (mean = 3.1 µm, sem 0.2, n=78; decrease = 1.6 µm; significant with p < 0.001) (Figure 5A). Interestingly, in *TgDCX* knockout parasites, fluorescently tagged TgCPH1 is targeted to the conoid, indicating that additional (non-tubulin) components must be involved in TgCPH1 targeting (Figure 5B). Together, these results indicate that TgDCX can influence the organization, curvature, and stability of the polymers, and bridge other conoid components to the tubulin core.

To determine which regions of the TgDCX molecule are critical for microtubule binding, bundling, and bending, we expressed in *Xenopus* S3 cells mCherryFP-tagged truncations of TgDCX that included the DCX domain only (aa 148-243), partial P25α domain only, or both the partial P25α domain and the DCX domain (aa 71-243), as well as domain-swapped versions containing 2 copies of the partial P25α, or 2 copies of the DCX domain, instead of one copy each. We found that neither mCherryFP tagged TgDCX148-243 (Figure 6A) (DCX-domain alone nor TgDCX135-256 (DCX domain plus part of linker region) (Supplementary Fig 2A) localizes to the microtubules, indicating that the DCX domain by itself is not sufficient for stable microtubule association. This agrees with the published data on human doublecortin binding to microtubules, where removing one of the two DCX domains eliminates microtubule binding (Taylor et al., 2000). Interestingly, mCherryFP-TgDCX16-134 (partial P25α domain plus part of linker region) does bind to microtubules (Supplementary Fig 2B), although with prominent unbound cytoplasmic and nuclear fractions. Perhaps surprisingly, a tandem dual-DCX construct, in which a DCX domain is substituted for the partial P25α domain in TgDCX, does not have detectable binding to microtubules *in vivo* (Supplementary Fig 3A). In contrast, the reciprocal domain swap, creating a tandem dual P25α molecule, binds quite efficiently to microtubules in *Xenopus* S3 cells (Supplementary Fig 3B), but it does not generate the short curved microtubule arcs induced by unmodified TgDCX.

**Figure 6.**
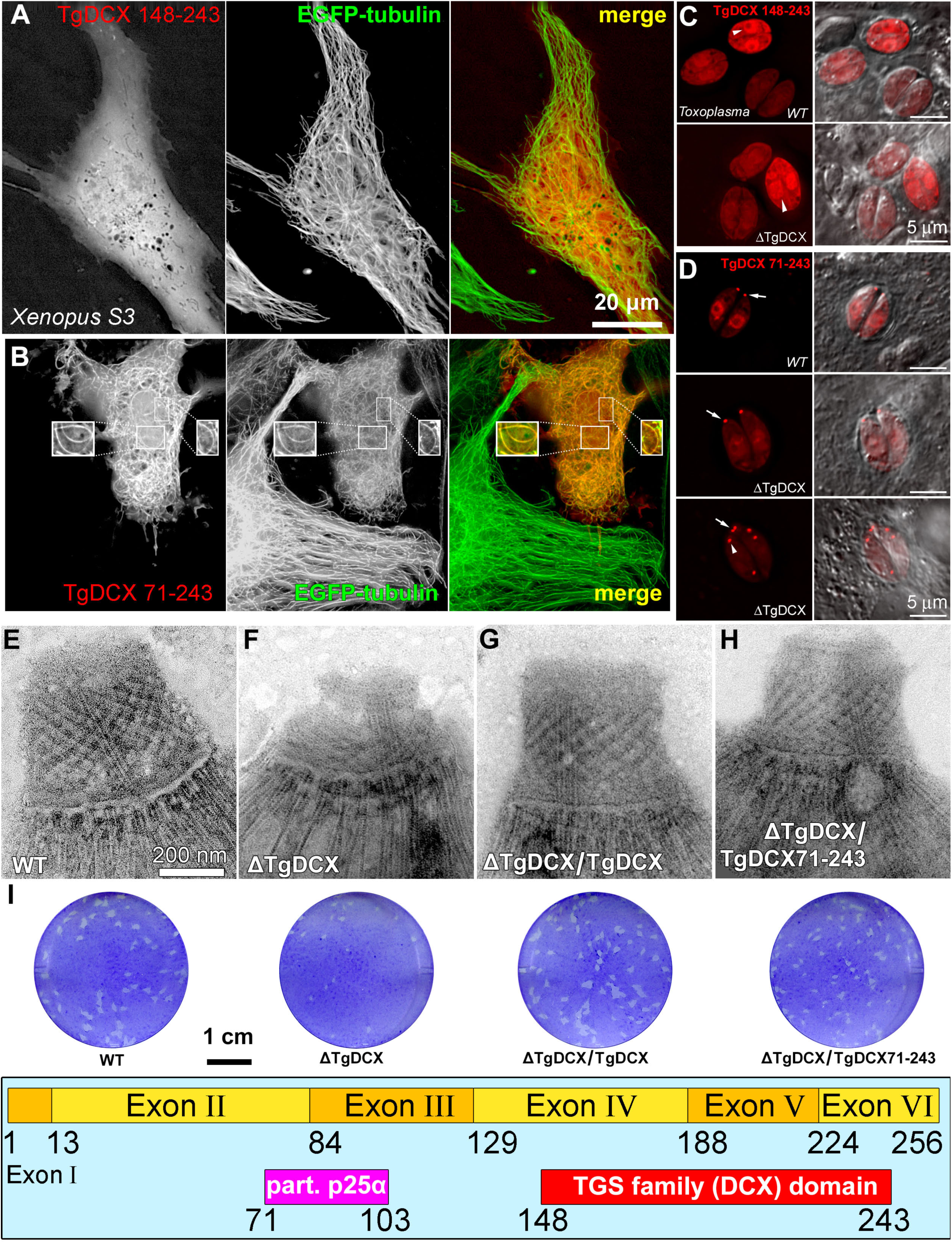
The DCX domain alone does not support stable microtubule binding in *Xenopus* cells, or conoid targeting in *Toxoplasma*. (A-B) Deconvolved wide-field images of *Xenopus* S3 cells expressing EGFP-tubulin (*green*) and either mCherryFP-TgDCX148-243 (A, *red*) or mCherryFP-TgDCX71-243 (B, *red*). The boxed insets in (B) are 1.5x enlarged and contrast-enhanced views of small regions over the nucleus that include the slices from the 3D stacks in which these individual arcs are clearly seen (also see Supplementary movie 3). The DCX domain alone (TgDCX148-243) is not sufficient for microtubule bending, but P25+DCX domain together (TgDCX71-243) cause bending to microtubules and generation of short arcs. (C-D) Deconvolved wide-field images of the parental RH*Δku80Δhx* (“WT”) and TgDCX knockout (“ΔTgDCX”) parasites expressing either mCherryFP-TgDCX148-243 (C), or mCherryFP-TgDCX71-243 (D, two examples are shown for ΔTgDCX). Arrowheads in (C) indicate the nucleus. Arrows in (D) point to the conoid; arrowhead in (D) points to a daughter conoid (E-H) EM images of the conoid region of negatively stained *T. gondii*. Parental RH*Δku80Δhx* (E, “WT”); *TgDCX* knockout (F, “ΔTgDCX”); knockout parasites transfected with a plasmid expressing either EGFP tagged full-length TgDCX (G, “ΔTgDCX / TgDCX”), or mCherryFP-TgDCX71-243 (H, “ΔTgDCX / TgDCX71-243”), both expressed under control of the *T. gondii* (constitutive, See Figure 9D) α-tubulin promoter. (I) Plaque assays (see *Methods*) of the parasite strains used for (E-H); the parental *T. gondii*, *TgDCX*-knockout parasites, and knockout parasites complemented with full-length TgDCX or the fragment containing only the partial P25α domain and the DCX domain, TgDCX71-243. Annotations are the same as (E-H). (J) Domain structure of TgDCX in which the amino acid boundaries of the partial P25α domain and the DCX domain are numbered.

Removing the N-terminal residues preceding the partial P25α domain of TgDCX has only a moderate effect. mCherryFP-TgDCX71-243 (partial P25α plus DCX domain) binds to and generates curved microtubule bundles in *Xenopus* S3 cells similar to those observed with the full-length protein, although with somewhat higher unbound cytoplasmic and nuclear fractions (Figure 6B, Supplementary movie 3). Thus, those 70 N-terminal aa contribute to binding efficiency, but are not required for microtubule binding, bundling, and bending. The difference in microtubule binding in *Xenopus* cells between constructs containing only the DCX domain (TgDCX148-243) versus constructs containing both the partial P25α and DCX domains (TgDCX71-243) is mirrored in conoid targeting in *Toxoplasma* (Figure 6C&D). When expressed in *Toxoplasma* (whether in the wild-type or in the TgDCX knockout background), mCherryFP-TgDCX148-243 was not enriched in the conoid. Interestingly, a slightly longer fragment, eGFP-TgDCX135-256 (“TgDCX C-half”) also has a large cytoplasmic pool, but does bind very weakly to the adult conoid and is significantly enriched on daughter conoids and cortical microtubules (Supplementary Fig 4). The fragment that contains both the partial P25α domain and DCX domain (*c.f.* domain map in Figure 6I), mCherryFP-TgDCX71-243, shows considerably more prominent binding to the conoids of both mother and daughter parasites with a lower concentration in the cytoplasm (Figure 6D). Furthermore, expression of mCherryFP-TgDCX71-243 in the *TgDCX* knockout parasite largely reverses the dramatic structural and lytic cycle defects caused by a lack of TgDCX (Figure 6E-I). In contrast, we were unable to obtain a transgenic line stably expressing mCherryFP-TgDCX148-243, indicating that transfection of *TgDCX* knockout parasites with this truncated form does not confer a growth advantage, possibly because conoid targeting is required for the function of TgDCX.

### Comparison of the structure, microtubule association, conoid targeting, and functional complementation between TgDCX and its orthologues

Organisms that contain TgDCX orthologues in their genomes differ greatly in lifestyles (*e.g.* obligate intracellular parasites *vs.* free-living phototrophs) and in conoid structures [*e.g.* a truncated cone made of non-tubular polymers in *Toxoplasma vs.* a sheet of canonical microtubules curved into an open-sided incomplete cone in *Chromera*, (Portman et al., 2014)]. Some of these organisms, such as *Trichoplax,* are thought to have no conoid-like structure.

The sequence conservation among the TgDCX orthologues is mostly restricted to the predicted P25-alpha and doublecortin domains (Figure 7A, Table 1) (Orosz, 2009; Orosz, 2016). Both the full-length TgDCX and the TgDCX71-243 fragment are insoluble in non-denaturing buffer systems (see *Methods*), but we were able to purify and crystallize a soluble fragment (AA148-243) that contains the DCX domain (Figure 7 and Supplementary Table 1) and determine its structure by X-ray crystallography. The fold of TgDCX148-243 is very similar to that of the N-terminal DCX domain of human doublecortin (Figure 7B), which itself is quite similar (root mean square distance (rmsd) of ∼ 2Å) to the doublecortin C-terminal DCX domain. [HsDCX-N and HsDCX-C, (Burger et al., 2016; Rufer et al., 2018)]. Comparing the three structures over the region in which they all align gives: TgDCX (<u>6B4A</u>, chain A, aa150-222) *vs.* HsDCX-N (<u>5IOI</u>, chain E, residues 51-128, rmsd 1.905 Å); TgDCX (<u>6B4A</u>, chain A, aa150-222) *vs.* HsDCX-C (<u>5IP4</u>, chain E, residues 178-252, rmsd 1.99 Å). Residues responsible for the four tubulin contact regions between HsDCX-N and tubulin identified by structural docking (Fourniol et al., 2010) are in most cases similar.

**Figure 7.**
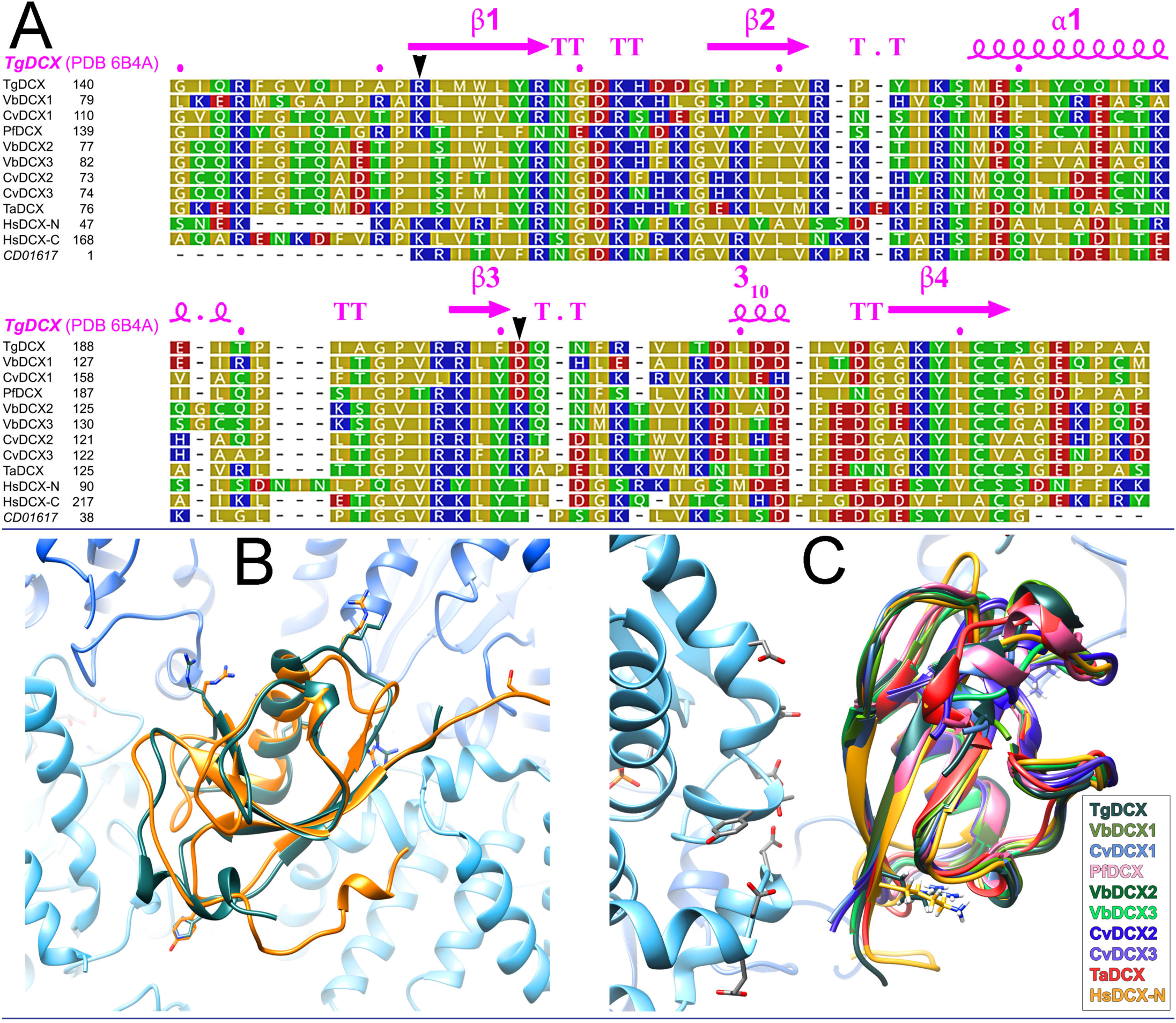
Sequence and structure conservation among DCX domains of TgDCX orthologues. (A) Sequences of DCX domains from *Toxoplasma gondii*, *Vitrella brassicaformis*, *Chromera velia*, *Trichoplax adhaerens*, N-and C-terminal domains of humans doublecortin, and a consensus DCX domain from the NCBI Conserved Domain Database (<u>CD01617</u>), were aligned using the MUSCLE program accessed through JalView (V2.10.5, http://www.jalview.org) with default parameters and displayed colored by polarity. Yellow: non-polar (G, A, V, L, I, F, W, M, P); Green: polar, uncharged (S, T, C, Y, N, Q); Red: polar, acidic (D, E); Blue: polar, basic (K, R, H). Secondary structure elements in TgDCX (beta sheet, “β”; alpha helix, “α”; turns, “T”; and a short stretch of 3_10_ helix), derived from our X-ray crystal structure (PDB <u>6B4A</u>) of TgDCX148-243, are indicated above the alignment. Black arrowheads indicate residues discussed in the text (TgDCX R152 and HsDCX-N K53; TgDCX D201). CvDCX1 (EupathDB ID: Cvel_6797), CvDCX2 (EupathDB ID: Cvel_18664), CvDCX3 (EupathDB ID: Cvel_28653), VbDCX1 (EupathDB ID: Vbra_15441), VbDCX2 (EupathDB ID: Vbra_12248), VbDCX3 (EupathDB ID: Vbra21191), PfDCX (EupathDB ID: PF3D7_0517800), TaDCX (Uniprot ID: B3RTF1) and HsDCX (NG_011750). (B) Superposition of backbone ribbon traces of TgDCX148-243 (dark gray-green; X-ray, <u>6B4A</u>) on the N-terminal DCX domain of human doublecortin (yellow-orange; solution NMR, <u>1MJD</u>, #11) docked with the structure of αβ-tubulin (α-tubulin blue, β-tubulin cyan; electron crystallography, <u>1JFF</u>) onto the cryoEM map of human doublecortin bound to microtubules (<u>4ATU</u>). The DCX domain binds in the groove between protofilaments, making contacts with two αβ-dimers. Side-chains are shown for some of the surface-exposed residues. The view is from the outside of the microtubule, corresponding to Figure 2C of (Fourniol et al., 2010). The (+)-end of the microtubule is towards the top. (C) Backbone ribbon threading of apicortin orthologue sequences onto the superimposed experimentally-determined structures of human DCX-N docked on microtubules (<u>4ATU</u>) and TgDCX148-243 (<u>6B4A</u>). The view is from the outside of the microtubule. The structures are rotated 135 degrees clockwise relative to the orientation shown in (B). The microtubule (+)-end points toward the bottom right corner of the diagram. Only the portion of one β-tubulin close to the DCX domain is included. Side chains are shown for the electronegative patch on β-tubulin, close to DCX, identified as contact region #3 on β-tubulin by (Fourniol et al., 2010). Side-chains of the residues discussed in the text, K53 on HsDCX-N and R152 on TgDCX are also shown, pointing away from tubulin in these structures.

**Table 1.**
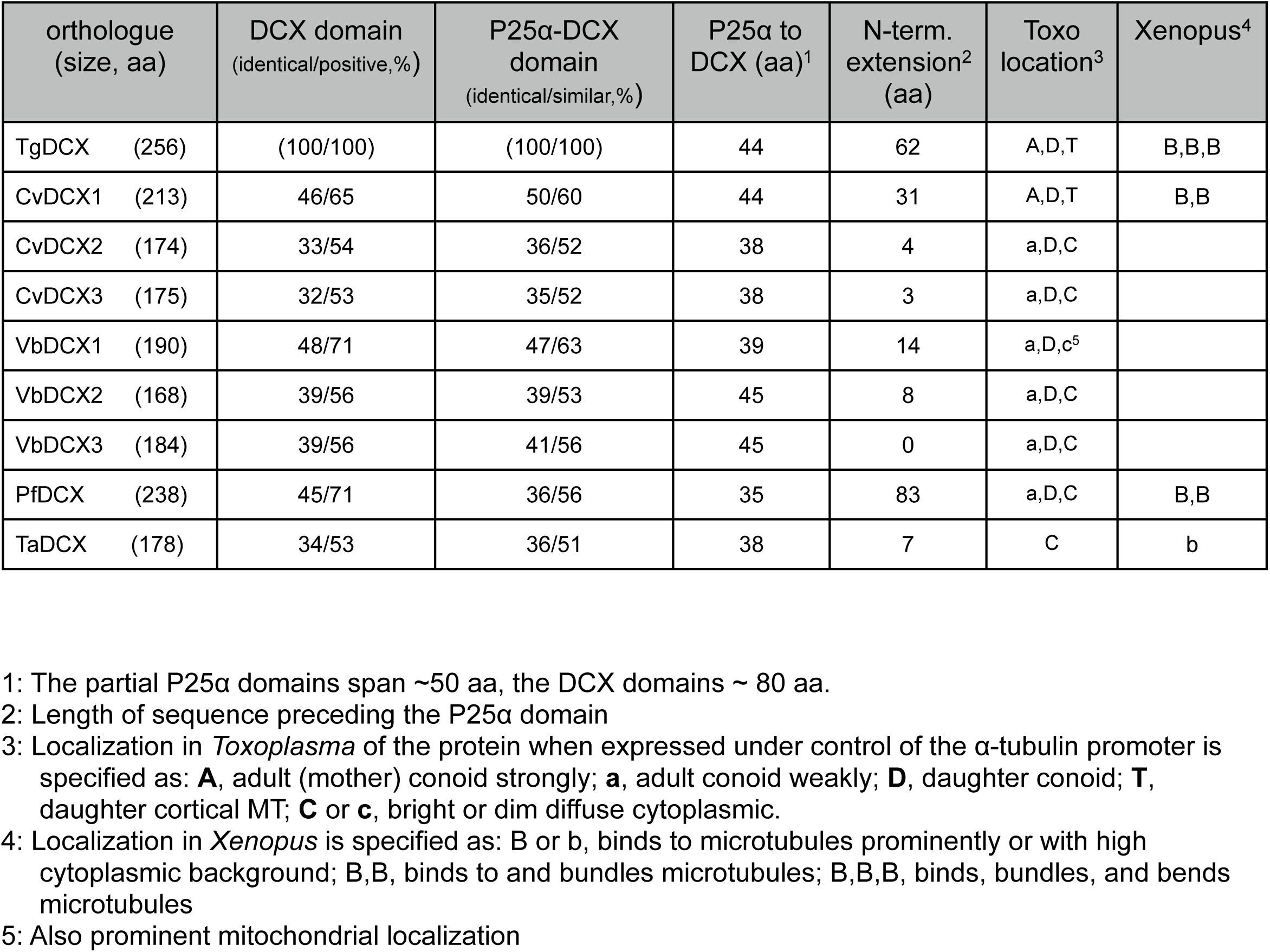
Sequence conservation and microtubule interaction properties among TgDCX and its orthologues.

Given the high level of homology among the TgDCX orthologues in the region of their DCX domains (Figure 7A, Table 1), we used the structure of TgDCX148-243 as a model for threading (Figure 7C) the corresponding regions in the TgDCX orthologues in *Chromera velia* (CvDCX1, CvDCX2, CvDCX3), *Vitrella brassicaformis* (VbDCX1, VbDCX2, VbDCX3), *Plasmodium falciparum* (PfDCX), and *Trichoplax adhaerens* (TaDCX). At the location reported in (Fourniol et al., 2010) corresponding to tubulin contact region #3 in HsDCX-N (K53), which abuts a strongly electronegative patch on β-tubulin, orthologues that bind efficiently to microtubules have a positive charge (R152 in TgDCX, lysine in CvDCX1 and PfDCX), whereas most of the orthologues that do not bind have isoleucine. The comparison of orthologues also revealed a reversal of polarity at one position: D201 in TgDCX and D at the corresponding location in CvDCX1 and PfDCX, is replaced by K or R at the corresponding residue in CvDCX2, CvDCX3, VbDCX2, VbDCX3, and TaDCX. Both sidechains however are orientated away from the microtubule, indicating that they are not directly involved in tubulin binding. Indeed, mutations at these two residues in TgDCX (R152I_D201R) and in a non-binding orthologue CvDCX2 (I85R_R134D) do not appreciably decrease TgDCX binding to microtubules nor increase CvDCX2 binding in *Xenopus* S3 Cells (Supplementary Fig 5).

To examine the extent of conservation in microtubule and conoid association among these orthologues, we expressed mCherryFP tagged orthologues of TgDCX in *Xenopus* S3 cells (Figure 8) and in *Toxoplasma* (Figure 9). No significant microtubule localization was found for mCherryFP tagged CvDCX2, CvDCX3, and the three VbDCX orthologues in *Xenopus* cells (Figure 8, Table 1). In contrast, CvDCX1 and PfDCX are prominently localized to the microtubules. Some mCherryFP-TaDCX associates with microtubules but the unbound cytoplasmic fraction is much larger relative to the bound fraction than is the case for the three orthologues that do bind strongly.

**Figure 8.**
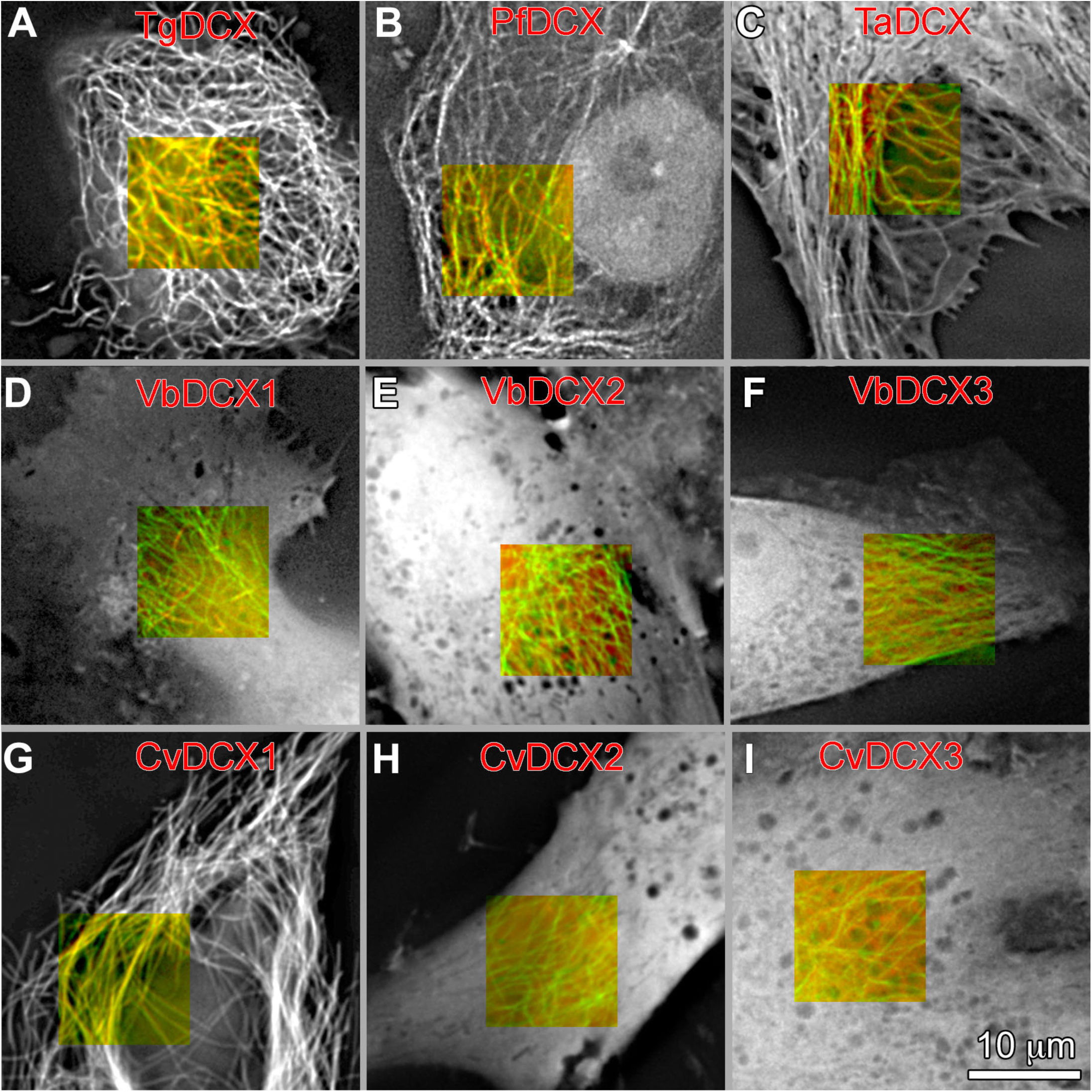
Microtubule binding in *Xenopus* S3 cells by TgDCX and its orthologues. (A-I) Deconvolved wide-field images of *Xenopus* S3 cells expressing both EGFP-tubulin and mCherryFP tagged DCX orthologues. The grayscale images show the fluorescence from the mCherryFP channel only. For a subarea, both channels are displayed to show both the EGFP tagged microtubules (*green*) and mCherry tagged orthologues (*red*). In A, B, C, and G, the orthologues are associated with microtubules, whereas in D, E, F, H, and I, they are distributed diffusely throughout the cytoplasm and nucleus.

**Figure 9.**
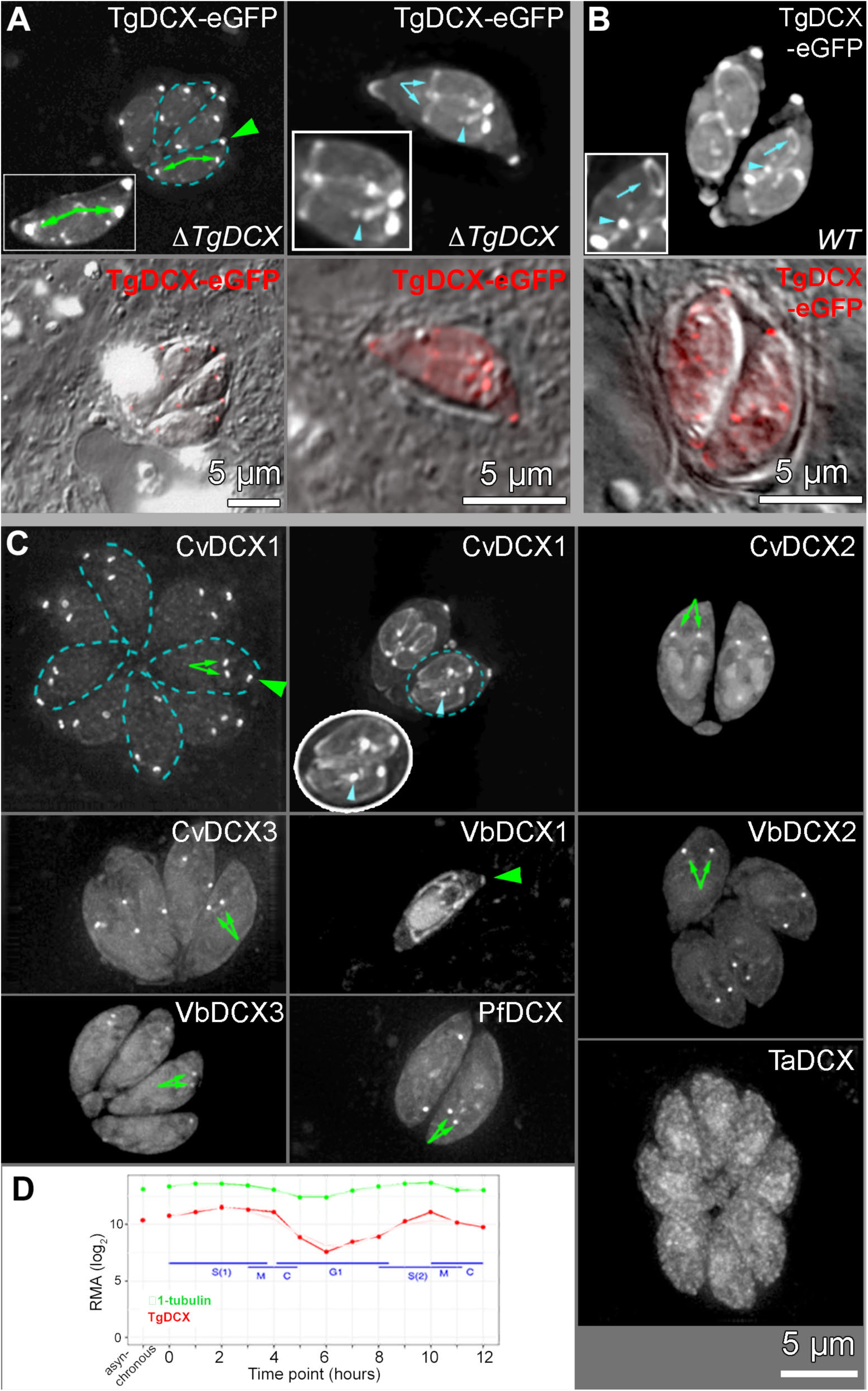
Localization of TgDCX and its orthologues in *Toxoplasma* (A-B) Deconvolved wide-field images of dividing *TgDCX* knockout (A, “ΔTgDCX”, two examples) and RH*Δku80Δhx* (B, “WT”) parasites transiently expressing TgDCX-eGFP driven by the *T. gondii* α-tubulin promoter. TgDCX-eGFP is highly enriched in the mother conoid (*green arrowhead*) and daughter conoids (*green arrows*) and is absent from the cortical microtubules of mother parasites. However, in contrast to expression regulated by the endogenous promoter, when expression is driven by this nearly constitutive (see D) α1-tubulin promoter, in some cases TgDCX-eGFP signal is also detected on the daughter cortical microtubules, centrosomes (*cyan arrowheads*), and basal complexes (*cyan arrows*). Dashed cyan lines in (A) outline two of four parasites in the same parasitophorous vacuole. Insets: 1.5x. The lower panels show merged DIC and fluorescence (in red) images. (C) Deconvolved wide-field images of RH*Δku80Δhx* (WT) parasites expressing FP tagged DCX orthologues. Two examples are shown for CvDCX1. In the left example, dashed blue lines outline 4 of the 8 parasites in the vacuole. In the right example, the dashed blue oval outlines two nearly mature daughters, shown enlarged 1.5x in the oval inset with white outline. Note that among the eight orthologues, only CvDCX1 closely mimics the pattern of localization shown by TgDCX (when expressed under this *T. gondii* α1-tubulin promoter). *Green arrows*: daughter conoids. *Green arrowheads*: mother conoids. *Cyan arrowhead*: centrosome. (D) Time course of RNA expression levels (Behnke et al., 2010) in *Toxoplasma gondii* for α1-tubulin (*green*) and TgDCX (*red*). Tubulin expression is nearly constitutive, whereas TgDCX varies by more than 30-fold across the cell-cycle.

In *Toxoplasma*, we expressed FP-tagged orthologues under the control of the α-tubulin promoter (Figure 9). We have previously shown that TgDCX-eGFP expressed from an α-tubulin promoter can complement the structural, invasion and lytic cycle defects of the *TgDCX* knockout parasite [(Nagayasu et al., 2016), also see Figure 6]. When transiently expressed from this promoter, TgDCX-eGFP is targeted to the conoid efficiently both in wild-type and *TgDCX* knockout parasites (Figure 9A-B). When expression is driven by the endogenous TgDCX promoter, both FP tagged [mCherryFP-TgDCX (Figure 1B-D), and TgDCX-mNeonGreenFP] and untagged TgDCX localize exclusively to the conoid (Nagayasu et al., 2016). In contrast, when expression is driven by the α-tubulin-promoter as in Figure 9A and 9B, TgDCX sometimes decorates the daughter cortical microtubules, centrosomes and basal complex. This reveals that the specificity of conoid association by the TgDCX protein is not absolute, and likely modulated by regulation at the transcription and post-translational level. Notably, while FP-tagged TgDCX can coat the daughter cortical microtubules when expressed from the α-tubulin promoter [which is essentially constitutive, (Behnke et al., 2010), Figure 9D], it does not stay bound to the cortical microtubules of the adult parasite, revealing a change in the surface property of the cortical microtubules as the parasite becomes mature.

The conoid targeting efficiency of the TgDCX orthologues varies widely (Figure 9C, Table 1). TaDCX is not enriched in the conoid. CvDCX2, CvDCX3, VbDCX2, VbDCX3, and PfDCX are prominently associated with the daughter conoid, but barely detectable in the mother conoid. This reveals that while conserved features in these orthologues allow them to interact with components of an assembling conoid, the structural tolerance for the divergent features decreases as the conoid matures. VbDCX1 is targeted to both mother and daughter conoids but also shows prominent unbound cytoplasmic, mitochondrial and nuclear pools. CvDCX1, which shares the highest sequence similarity with TgDCX (44% identical, 57% similar over the full-length), largely mimics the localization of TgDCX when its expression is driven by the α-tubulin promoter. It is prominently localized to the conoids of both adult and daughter parasites, and sometimes to the cortical microtubules and centrosomes of daughter parasites, with only a small cytoplasmic pool.

Despite the similar localization pattern between CvDCX1 and TgDCX, when CvDCX1 is expressed in the *TgDCX* knockout background, it fails to complement the defects in conoid structure and in the lytic cycle (Figure 10). EM examination of negatively-stained whole-mount parasites showed that *ΔTgDCX* parasites expressing CvDCX1 still have a distorted conoid structure like the knockout parasites. Contrast this with the restored conoid structure in *ΔTgDCX* parasites expressing full-length TgDCX or TgDCX71-243 (Figure 6G and H).

**Figure 10.**
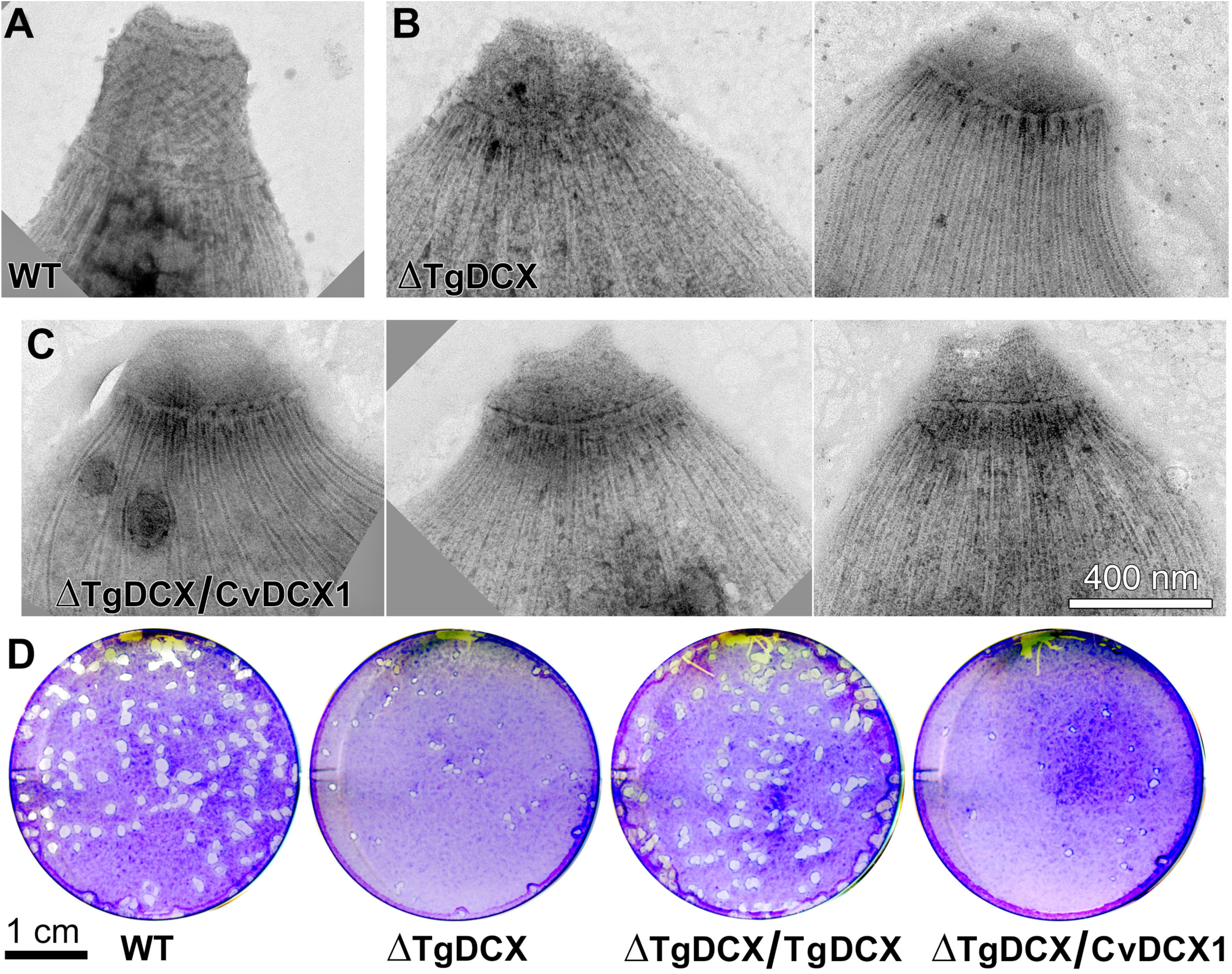
CvDCX1 cannot rescue the structural and lytic cycle defects of the TgDCX knockout parasite. (A-C) EM images of the conoid region of negatively stained *T. gondii*. Parental RH*Δku80Δhx* (A, “WT”), *TgDCX* knockout (B, “ΔTgDCX”, two examples); a clone of *TgDCX* knockout parasite stably expressing CvDCX1-mNeonGreenFP under control of the *T. gondii* α-tubulin promoter (C, “ΔTgDCX / C vDCX1”, three examples). Compare with Figure 6G and H. (D) Plaque assay (see *Methods*). Knockout parasites complemented with TgDCX-eGFP expressed under control of the *T. gondii* α-tubulin promoter (“ΔTgDCX / TgDCX “). Other annotations as in (A-C). Compare with Figure 6I.

To further determine how TgDCX and CvDCX1 differentially modify the organization and structure of tubulin polymers, we carried out correlative light and electron microscopy analysis of *Xenopus* S3 cells expressing FP-tagged TgDCX, TgDCX71-243, and CvDCX1 (Figure 11 & 12). In untransfected (Figure 11A and B) and FP-CvDCX1 transfected (Figure 11C-E) *Xenopus* S3 cells, microtubules invariably have 13 protofilaments (pf) and are closed tubes. In contrast, TgDCX (Figure 12A) or TgDCX71-243 (Figure 12B) containing fibers are mostly single-layered, occasionally multi-layered, rafts of microtubules, each layer containing 2-10 microtubules arranged side by side in a plane. The planar raft-like arrangement is most simply understood as the natural consequence of “zipping” curved microtubules together: the distance over which neighboring microtubules remain in contact is greatly increased if the microtubules are stacked side-by-side, all bending in the same direction. Notably, ∼40% of the rafts are bordered on one edge by an incomplete tube. This “C”-shaped element appears to be a 13-protofilament “microtubule” with an opening on one side, the side that faces the edge of the sheet. The “C”-shaped incomplete tubes are also seen as single fibers, not associated with a raft. Among all the microtubules with clear profiles in three different cells, 36% were “C”-shaped incomplete tubes (114 out of 313). The diameters of TgDCX or TgDCX71-243 containing tubes are often greater than the canonical 13-pf microtubules. Some of those microtubules contain more than 14 protofilaments (Figure 12C).

**Figure 11.**
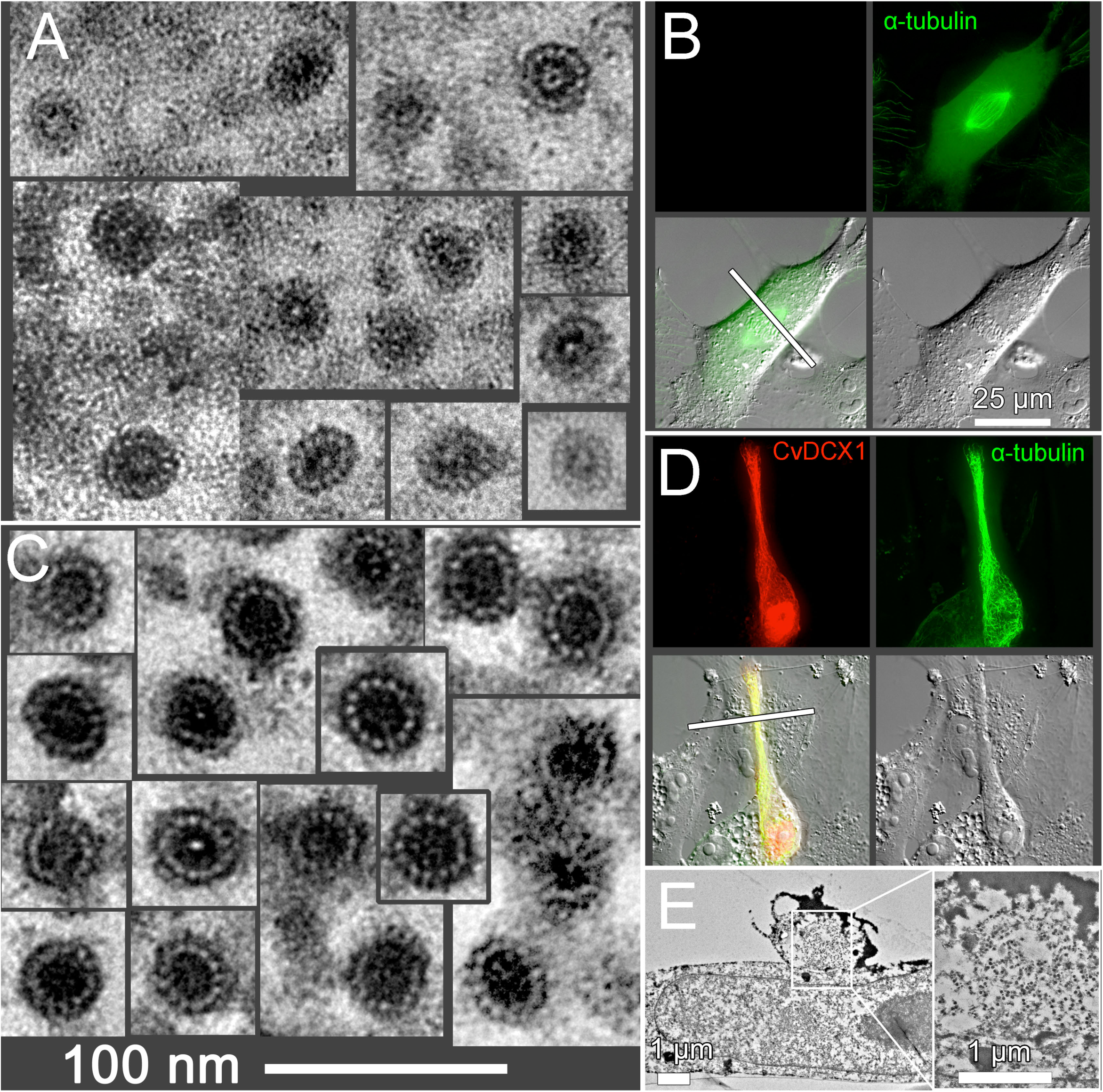
Correlative light and electron microscopy analysis of microtubules in untransfected CvDCX1-expressing *Xenopus* cells. (A) EM images of microtubules in sections of an untransfected *Xenopus* cell. In all cases where they are countable, 13 protofilaments (pf) are present. No microtubules with other than 13 pf were seen in untransfected cells. (B) Fluorescence and DIC light microscope images of the cell sectioned in (A). The cell is from a line expressing EGFP-α-tubulin. The plane of sectioning in the EM images is shown by the white bar. (C) EM images of microtubules in sections of a *Xenopus* cell transfected with mCherryFP-CvDCX1. All microtubules have 13 pf. The tannic-acid-enhancement of microtubule staining (see *Methods*) is more effective when the microtubules are heavily decorated, which makes the protofilaments more obvious and more easily countable, but the diameter of the microtubules is approximately the same as in untransfected cells. (D) Fluorescence and DIC light microscopy images of the cell sectioned in (C). mCherryFP-CvDCX1 is shown in red, EGFP-α-tubulin in green. The plane of section is shown by the white bar. The magnification is the same as for (B). Note that the elongated narrow extension of the transfected cell lies on top of another untransfected cell, running over the edge of the latter’s nucleus. (E) Low magnification EM images of a cross-section of the cell shown in D. The thin extension of the transfected cell is seen crossing over the underlying untransfected cell. The region within the white box, shown enlarged on the right, contains >100 parallel microtubules viewed in cross-section, appearing as tiny black doughnuts at this magnification.

**Figure 12.**
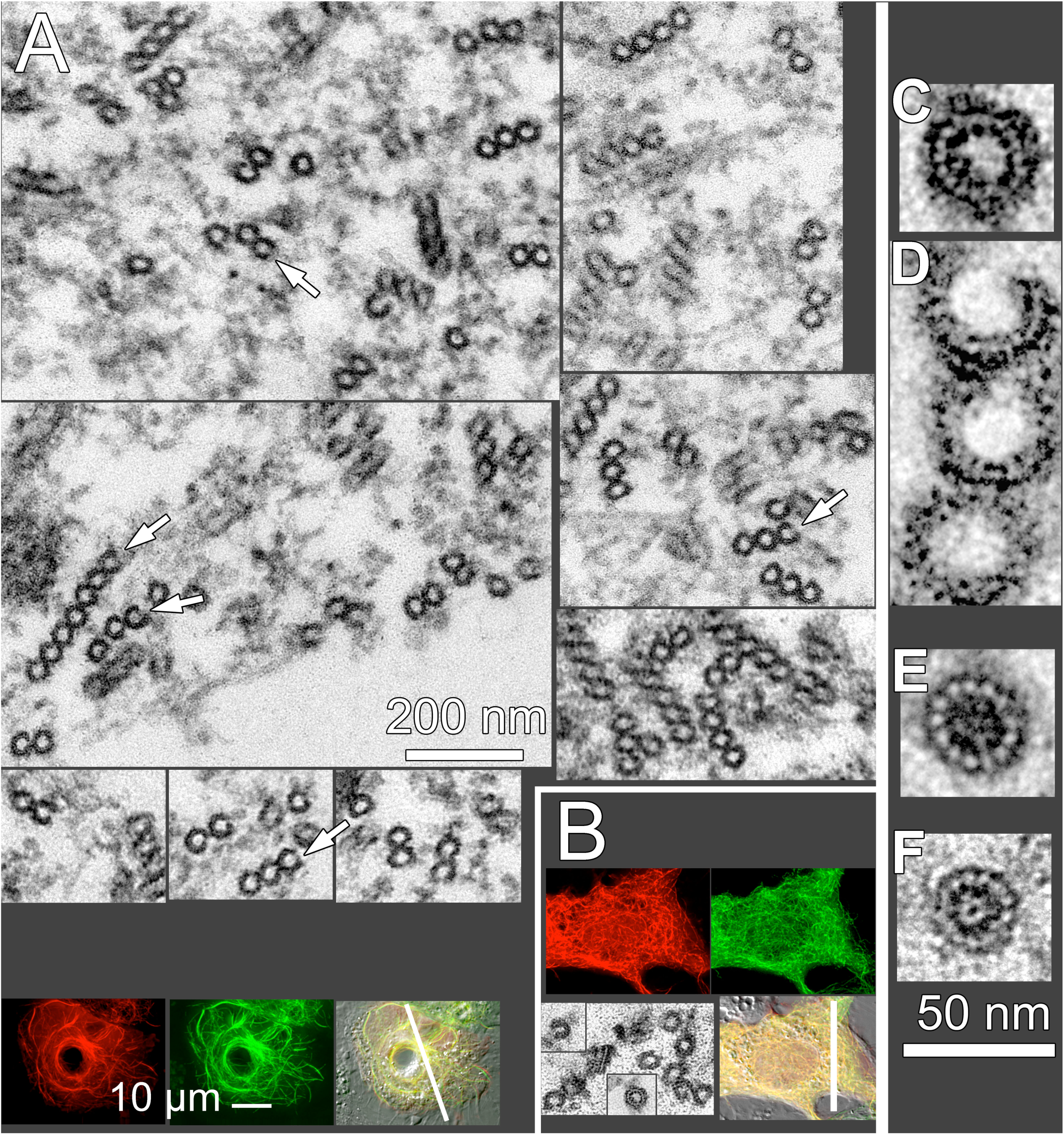
Correlative light and electron microscopy analysis of microtubules in *Xenopus* cells expressing TgDCX. (A) Montage of images of microtubule rafts viewed in cross-section by EM, from a *Xenopus* cell expressing TgDCX-mCherryFP. Often the microtubules on one edge of a raft are incomplete tubes (*white arrows*), as are some of the single microtubules in these cells. The light micrographs at the bottom show fluorescence and DIC images of the sectioned cell, which is also expressing EGFP-α-tubulin. The plane of sectioning is shown by the white bar. TgDCX-mCherryFP is shown in red, EGFP-α-tubulin in green. (B) Light and electron microscope images of a *Xenopus* cell expressing mCherryFP-TgDCX71-243 and EGFP-α-tubulin. mCherryFP-TgDCX71-243 is shown in red, EGFP-α-tubulin in green. The plane of section is indicated by the white bar. The EM images show microtubule cross-sections in these cells, which are similar to those in cells expressing full-length TgDCX, quite different from microtubules in untransfected cells (*c.f.* Figure 11). (C-F) Comparison of microtubule shapes and sizes. C&D: a single microtubule with more than 13 pf, and a cluster of three microtubules from TgDCX71-243 transfected cells. Note that the width of the incomplete microtubules is often larger than the complete tubes, suggesting that the gap results from a tube expanding in diameter and splitting open, rather than from loss of protofilaments. E: a 13 pf microtubule from a cell expressing CvDCX1. F: a 13 pf microtubule from an untransfected cell.

Thus, while the sequence conservation between CvDCX1 and TgDCX is high and sufficient for similarly specific localization to the conoid, enough variations have accumulated during the evolutionary divergence of these two lineages to result in a difference in their abilities to restore the conoid structure of the *TgDCX* knockout parasite and support the parasite lytic cycle. This is likely related to a difference in their impact on the arrangement of the protofilaments, and on the organization and curvature of the tubulin polymers.

## DISCUSSION

The conoid originated early in the evolutionary history of the apicomplexan parasites. It is present in all coccidians examined to date (*e.g. Toxoplasma* and *Eimeria* spp.). Although apicomplexans in *Haemosporidia* (*e.g. Plasmodium* spp.) are classified as *Aconoidasida* (“conoidless”), a conoid-like structure was found in the ookinete stage of *Plasmodium* and other haemosporidians by EM (Desser et al., 1970; Patra and Vinetz, 2012). A conoid-associated protein (SAS6L) was also found in the sporozoite and ookinete stages of *Plasmodium* highlighting a similar apical structure as its homolog in *T. gondii* (Wall et al., 2016). The level of *PfDCX* transcription relative to the whole transcriptome is much higher in the gametocyte and ookinete stages compared with those in the blood stages. Furthermore, even though it has a divergent P25α domain, PfDCX binds strongly to microtubules when expressed in *Xenopus* S3 cells and to daughter conoids when expressed in *Toxoplasma.* It is therefore a useful probe to determine whether *Plasmodium* spp assemble a conoid-like structure at certain stages of their life-cycle, and the organization of the protofilaments in the “conoid” fibers if such structure exists.

It is interesting to consider the geometry of the arrangement of the fibers in the conoid, particularly with respect to the changes that must occur upon conoid extension (*c.f.* the model in Figure 2E). In the retracted state, the conoid is a truncated cone, widening from a diameter of ∼400 nm at the top to ∼600 nm at its basal end, with conoid fibers angled at ∼25° relative to base of the conoid (Hu et al., 2002). For this geometry, the conoid fibers follow a path with radius of curvature varying from 250 to 260 nm, close to what is observed for isolated fibers in the cryoEM images. After extension, the apical diameter is unchanged, but the basal diameter has decreased to ∼450nm, the conoid has become taller and more cylindrical, and the angle of the conoid fibers has increased to ∼40°. The fibers then follow a more steeply inclined, and therefore less curved path: the radius of curvature for the fiber path in the extended conoid varies from ∼300 to ∼340 nm, somewhat larger than observed for isolated fibers. From these admittedly approximate calculations, we propose that when released from the conoid, the fibers adopt the configuration characteristic of the retracted state, and that this therefore is the lower energy, most strain-free, configuration. The change in pitch during conoid extension results in the fibers becoming more parallel to the conoid central axis. Though the consequences of this change in pitch are not immediately obvious (at least, were not obvious to the authors), carefully thinking about this geometry reveals that the change in pitch necessitates a change in twist of the fiber (unless both ends are free to rotate, which seems less plausible). When the conoid extends, the diameter at the base decreases, with concurrent clockwise (viewed from the apical end) rotation of the base with respect to the apex, thus partially “unwinding” the left-handed spiral of the conoid fibers. The consequence of this “unwinding” for fibers anchored at their ends, as the conoid fibers seem to be, is twisting along the length of each individual fiber in a right-handed manner. However, sagittal sections of the conoid in intact parasites (Hu et al., 2002) show that in the extended state, the conoid fibers all have the same rotational orientation (right half of the model in Figure 2E). A sagittal section cuts each fiber at a different location along its length, from closer to their apical end for the fibers at the top of the section, to near the basal end for fibers at the bottom of the section. Thus, the observed identical rotational orientations of the fibers at the top and bottom demonstrate that the fibers have little or no twist in the extended conoid. This implies that they had a left-handed twist in the retracted conoid (left half of the model in Figure 2E), and then untwisted as a consequence of conoid extension. This again suggests that the fibers seen in the cryoEM images (*e.g.* Figure 2A) are in the configuration (*i.e*., twisted) they have in the retracted conoid.

The apicomplexan parasites are thought to have evolved from a free-living marine protozoan with conserved ancestral traits and cellular features. Indeed, a “pseudoconoid” is present in the marine relatives of the apicomplexans, such as *C. velia* and *V. brassicaformis.* In *Chromera*, the pseudoconoid is a half-cone formed from a sheet of 30-35 canonical microtubules (Portman et al., 2014), in which the curvature of the microtubules is much less than that of the conoid fibers in *Toxoplasma.* Thus, the conoid underwent considerable structural rearrangement during the evolution of the intracellular parasitic lifestyle of the apicomplexans. Characterization of the components of the conoid might help to understand the molecular basis of these lifestyle transitions. If we presume that the pseudoconoid as seen in *Chromera* is similar to the ancestral form of the conoid in *Toxoplasma,* then the ancestral form of TgDCX very likely binds to microtubules, which form the pseudoconoid. Our observation that TgDCX can associate with microtubules in *Xenopus* S3 cells is consistent with this hypothesis. However, as TgDCX binds quite readily to canonical microtubules, it is puzzling that it can be specifically localized to the conoid in *Toxoplasma* in the presence of several other tubulin-containing structures (Figure 1, *mCherryFP-TgDCX* knock-in parasite), particularly puzzling since the identical tubulin subunits are assembled into the non-tubular conoid fibers simultaneously and in close proximity to assembly into canonical (tubular) microtubules during daughter construction (Figure 1C-D) (Nagayasu et al., 2016). When expressed from a strong constitutive promoter, FP-tagged TgDCX is predominantly localized to the conoid, suggesting preferential binding to non-tubular tubulin polymers (Figure 9). However, expressed in this manner, TgDCX also binds to daughter cortical microtubules, indicating that the specific conoid targeting in *Toxoplasma* must involve other levels of control, such as transcriptional regulation in combination with post-translational modifications of TgDCX and its targets.

Similar to TgDCX, CvDCX1 from *Chromera* was observed to associate with microtubules in *Xenopus* S3 cells and be targeted to the conoid in *Toxoplasma*. However, given the strong sequence conservation of the TgDCX orthologues in the partial P25α domain and the DCX domain, including the residues predicted to serve as tubulin contacts, it is surprising that the efficiency of microtubule and conoid association vary so widely among the other orthologues. For instance, all three orthologues in *Chromera* share a high degree of conservation with TgDCX, but CvDCX2 and CvDCX3 show no significant enrichment on the microtubules in *Xenopus* S3 cells and are not efficiently targeted to the conoid of mature parasites when expressed in *Toxoplasma.* While we predict that CvDCX1 is localized to the pseudoconoid in its native environment, it will be of interest to characterize all three orthologues in *Chromera* and determine whether they have overlapping or distinct localization and functions.

During the divergence of the apicomplexans, the precursor gene for *TgDCX* must have accumulated *Toxoplasma-*specific features for maintaining the structural integrity of the conoid and facilitating invasion, as the expression of the orthologue (CvDCX1) that most closely mimics the localization of TgDCX in *Toxoplasma* fails to restore the conoid structure and also fails to complement the lytic defects of *TgDCX* knockout parasite. One such specific feature might be TgDCX’s ability to generate and stabilize curved tubulin polymers (Figures 3-4), which might contribute to the stabilization of the highly curved conoid fibers of *Toxoplasma.* However, we note that TgDCX is unlikely to be the only factor that contributes to the bending of the conoid fibers, because: 1) distorted curved fibers can sometimes be observed in the conoid of the TgDCX knockout parasite (Figure 2); 2), the radius of curvature of the arced microtubules seen in *Xenopus* cells is significantly greater than that of the conoid fibers (4.7 *vs*. 0.25 µm); and 3), the incomplete tubes formed in *Xenopus* cells in the presence of TgDCX are more similar to microtubules than to the conoid fibers (in cross-section, incomplete circles rather than inverted “J” shapes). The co-expression of another conoid component, TgCPH1, slightly decreases the radius of curvature to ∼ 3.1 µm, which is still more than an order of magnitude larger than that of the conoid fibers.

Interaction with microtubules differs dramatically between TgDCX and its orthologues, also likely differs between TgDCX and human doublecortin. In the case of doublecortin, a detailed model of the interaction of its two DCX domains (NDC and CDC) with microtubules has been proposed, based on high-resolution structural information from cryoEM (Manka and Moores, 2019). A striking feature of the model is that for the most part, binding of the two domains to the microtubule is mutually exclusive: either NDC binds or CDC binds, with differing consequences for microtubule architecture, but apparently the two domains rarely or never bind simultaneously. This degree of independence seems unlikely to hold true for the two MT-binding domains of TgDCX. Full length or TgDCX71-243, which includes both P25α and DCX domains, induce dramatically different microtubule architecture compared to constructs containing just one of the two domains, or to constructs containing two copies of the same domain. It is difficult to imagine how such different architectures could come about unless both the P25α and the DCX domains interact simultaneously with a microtubule. Similar to TgDCX-N-half, which contains the P25α domain only, the *Chromera* and *Plasmodia* orthologues of TgDCX that do bind to microtubules in *Xenopus* cells do not generate short, curved microtubules. One possible explanation is that in these orthologues, the association with the MT is mediated predominantly by only a single domain, much as postulated for human doublecortin.

By combining localization, structural and cross-complementation analyses, we discovered that TgDCX orthologues from organisms with drastically different lifestyles have distinct polymer binding and modifying capabilities. In the future, systematic comparison of orthologues of conoid components in the apicomplexans and their marine relatives will be informative to determine if and how the molecular and structural changes in the conoid-like structures contribute to the transition from the free-living to the parasitic lifestyle of the apicomplexans.

## METHODS

### Culture, harvest, and transfection of Toxoplasma gondii

*T. gondii* tachyzoites were used in all experiments, and grown in monolayers of human foreskin fibroblast (HFF) cells (Roos et al., 1994). Transfection of *T. gondii* tachyzoites was carried out as previously described (Heaslip et al., 2010) using 30-40 µg of plasmid DNA in “cytomix” buffer (120 mM KCl; 0.15 mM CaCl_2_; 10 mM KH_2_PO_4_ / K_2_HPO_4_; 25 mM HEPES; 2 mM K_2_EGTA; 5 mM MgCl_2_, 2 mM K_2_ATP, 5 mM glutathione; pH adjusted to 7.6 with KOH).

### Plaque assay

Plaque assays were performed as previously described (Nagayasu et al., 2016).

### Xenopus S3 cell culture, transfections, and nocodazole treatment

Xenopus S3 cells stably expressing eGFP-tubulin (a kind gift from Drs. John Daum and Gary Gorbsky, Oklahoma Medical Research Foundation, Oklahoma City, OK) were maintained at room temperature (22°C) in 70% Leibovitz’s L-15 medium (ThermoFisher Scientific-Gibco, Cat#11415-064), 15% Fetal Bovine Serum, 15% H_2_O, 1% Penicillin/Streptomycin (ThermoFisher Scientific-Gibco, Cat#15140-122), and 72 μM G418 (ThermoFisher Scientific, Cat#10131035). For transfection, Lipofectamine® LTX Reagent with PLUS® Reagent (ThermoFisher Scientific, Cat#15338100) was used with the conditions recommended by the manufacturer, and 2.5 µg of plasmid DNA. In some experiments, ∼16 hours after transfection, cells were treated with 16.6 µM nocodazole (Sigma # M1404) for 30 min to 2 hours before imaging.

*Plasmid construction* (All PCR primers and synthesized gBlock gene fragments are listed in Supplementary Table 2).

After construction, plasmids were used to transform chemically competent TOP10 cells by heat shock, or electrocompetent DH5α cells (New England Biolabs, Cat# C2989) by electroporation. Plasmid DNA was isolated by standard procedures and the constructions were verified by DNA sequencing.

Plasmid ptub-EGFP-TgDCX was constructed on a ptub-EGFP-EGFP backbone, which has the same basic structure as ptub-H2b-YFP described in (Nagayasu et al., 2016), *i.e*., *Nhe*I-EGFP-*Bgl*II-EGFP-*Afl*II replaces *Nhe*I-H2b-*Bgl*II-YFP-*Afl*II, with the same 5 aa linker (SGRLS) separating the two inserts. Plasmid ptub-EGFP-TgDCX was constructed by cutting ptub-EGFP-EGFP with *Bgl*II and *Afl*II, and ligating in the *Bgl*II-*Afl*II digested TgDCX sequence PCR amplified from *Toxoplasma* cDNA using primer pair S19-AS19. To generate plasmids ptub-EGFP-TgDCX-N-half and ptub-EGFP-TgDCX-C-half, the TgDCX coding sequence was excised from ptub-EGFP-TgDCX with *Bgl*II-*Afl*II and replaced with *Bgl*II-*Afl*II cut PCR fragments of TgDCX (“N-half”, aa M16-V134, containing the partial P25-alpha domain, and “C-half”, aa T135-V256, containing the DCX domain plus linker region) amplified from *T. gondii* cDNA with primer pairs S19-AS24 and S24-AS19 respectively.

For the ptub-based plasmids driving expression in *Toxoplasma* of *Chromera* and *Vitrella* orthologues CvDCX1-3 and VbDCX1-3, ptubg-[DCX orthologue]-mNeonGreenFP was generated with a three-component assembly using the NEBuilder HiFi Assembly kit (New England Biolabs, E2621S) according to the manufacturer’s instructions. The vector backbone was prepared by removing the 12-base pair stuffer sequence in ptubg (Nagayasu et al., 2016) using *Nhe*I and *Afl*II. The mNeonGreenFP coding sequence was amplified using primers S1 and AS1 with the plasmid pmNeonGreenFP-N1 as a template [(Shaner et al., 2004; Shaner et al., 2013); a kind gift from Richard Day]. The coding sequence for the DCX orthologues including the ATG initiation codon but lacking the stop codon, and flanked by the linker sequences 5’-gaattcccttttagatccgctagcaaa-3’ at the 5’ end and 5’-cagatcctgttcctcc-3’ at the 3’ end, were synthesized as a gBlock gene fragment (Integrated DNA Technologies; Coralville, IA). To construct ptubg-TgCPH1-mCherryFP, a synthesized gBlock for TgGT1_266630 (“TgCPH1”) was used as PCR template with primer pair S23-AS23 and the product was combined with mCherryFP amplified with primer pair S1-AS1. Coding sequences for DCX orthologues of *Chromera velia* reference strain CCMP2878 and *Vitrella brassicaformis* reference strain CCMP3155 were obtained from CryptoDB (Heiges et al., 2006) (http://cryptodb.org/cryptodb/, release 41). Sequences for all gBlock gene fragments are listed in Supplementary Table 2.

ptubg_PfDCX-mNeonGreenFP was generated with a four-component HiFi assembly as described above for the *Chromera* and *Vitrella* orthologues, except Exons 1 and 2 of PfDCX were amplified using primer pairs S2 and AS2, and S3 and AS3, respectively, using *P. falciparum* 3D7A genomic DNA (BEI Resources, Manassas, VA, Cat# MRA-151G, contributed by David Walliker) as the template.

ptubg_TaDCX-mNeonGreenFP was generated with a three-component HiFi assembly as described above for the *Chromera* and *Vitrella* orthologues, except the coding sequence of TaDCX was amplified using primers S4 and AS4 using ptub-mEmeraldFP-TaDCX (synthesized by GenScript Inc, NJ) as the template.

To generate ptubg_mCherryFP-TgDCX148-243 or TgDCX71-243, the coding sequences for TgDCX truncations (TgDCX148-243 and TgDCX71-243) were PCR amplified from pTKO4_mCherryFP_TgDCX (Nagayasu et al., 2016) with primer pairs S5-AS5 and S6-AS6 respectively, and cloned into ptubg, fused to the C-terminus of mCherryFP (PCR amplified with primers S7-AS7), in a three-component NEBuilder HiFi assembly as described above for the TgDCX orthologues.

To generate mCherry tagged full-length wild-type TgDCX, full-length TgDCX-R152I-D201R, and the two TgDCX truncations for expression in *Xenopus* cells, the coding sequences were PCR amplified with primer pairs S20-AS20 for full-length, S8-AS8, and S9-AS9 respectively (with template pTKO4_mCherryFP_TgDCX for wild-type TgDCX, and synthesized TgDCX-R152I-D201R gBlock for the mutant) for cloning into an expression plasmid utilizing the CMV promoter (“pC22”). pC22 was derived from Clontech vector pEGFP-C2 by cutting with *EcoR*I plus *Xma*I, and inserting the adaptor formed by hybridization of oligos S10 and AS10. An intermediate plasmid, pC22_mCherryFP-HsTUBA1, was constructed from the *Nhe*I-*Afl*II fragment of pC22 by ligation of a PCR fragment containing mCherryFP-HsTUBA1, amplified with primers S11 and AS11 from plasmid pCMV_mCherryFP_HsTUBA1, a gift from the laboratory of Dr. Roger Tsien (Shaner et al., 2004). The human tubulin sequence in pC22_mCherryFP-HsTUBA1 was then excised with *Bgl*II-*Afl*II and replaced with the PCR amplified TgDCX fragments via a two-component NEBuilder HiFi assembly reaction. Plasmid pC22-TgDCX-EGFP was assembled into the *Nhe*I-*Afl*II-cut pC22 vector backbone by ligating in TgDCX-EGFP excised with *Nhe*I-*Afl*II from plasmid ptub-TgDCX-EGFP [described in (Nagayasu et al., 2016)]. Removing EGFP with *Nhe*I-*Bgl*II and replacing it via NEB HiFi assembly with mCherryFP amplified using primer pair S22-AS22 yielded plasmid pC22-TgDCX-mCherryFP. The “N-half” and “C-half” *BglII-AflII* fragments of plasmids ptub-EGFP-TgDCX-N-half and ptub-EGFP-TgDCX-C-half were ligated into *BglII-AflII* cut pC22_mCherryFP-HsTUBA1 to give the corresponding CMV-promoter driven expression plasmids.

Expression plasmids for domain-swapped versions of TgDCX were constructed for *Xenopus* (pC22_mCherryFP_DCX-DCX and pC22_mCherryFP_P25-P25) by NEBuilder HiFi assembly from synthesized gBlock fragments and *Bgl*II-*Afl*II cut vector backbones as described for ptubg and pC22 constructs above. The gBlock for the double-DCX version coded for aa M1-P62, G145-K248, H119-F144, and G145-V256 of TgDCX. To simplify cloning, sequencing, and PCR analysis, the nucleotide sequence corresponding to the first copy of the DCX domain, G145-K248, was comprehensively mutated at the third nucleotide of every codon (except ATG), as well as all three nucleotides for arginine, leucine, and serine codons, resulting in a 100% identical amino acid sequence but only ∼60% nucleotide sequence identity. The gBlock for the double-P25 version coded for aa M1-F144, A63-E118, and V249-V256 of TgDCX. The second copy of the P25 region, A63-E118, was mutagenized in the same way as for the DCX-DCX gBlock.

As described above for pC22-mCherryFP-TgDCX, CMV-promoter expression plasmids for the TgDCX orthologues were similarly derived from pC22_mCherryFP-HsTUBA1 using orthologue inserts PCR-amplified with primers S12-AS12 (Cvel_6797), S13-AS13 (Cvel_18664), S14-AS14 (Cvel_28653), S15-AS15 (Vbra_12284), S16-AS16 (Vbra_15441), S17-AS17 (Vbra_21191), and S18-AS18 (Pf3D7_0517800). A plasmid expressing a mutated version of Cvel_18664 (Cvel_18664_I85R-R134D) was similarly constructed using synthesized Cvel_18664_I85R-R134D gBlock as template for PCR. A pC22 expression plasmid for mCherryFP-TgGT1_266630 (“TgCPH1”) was constructed by PCR amplifying TgCPH1 from a synthesized gBlock with primer pair S21-AS21, and assembling it into *Bgl*II-*Afl*II-cut pC22-mCherryFP-TgDCX via a two-component NEBuilder HiFi assembly reaction. pC22-mCerulean3FP-TgCPH1 and pC22-mNeonGreenFP-TgCPH1 were derived from pC22-mCherryFP-TgCPH1 by excising mCherryFP with *Nhe*I-*Bgl*II and replacing it with one of the other FP’s PCR amplified with primer pair S22-AS22.

To construct the plasmid for expressing His-tagged recombinant TgDCX148-243 (BG1861_ TgDCX148-243), the coding sequence was amplified from the genomic DNA of *Toxoplasma gondii* strain ME49 and cloned into an *E. coli* expression vector (BG1861) using ligase-independent cloning (Aslanidis and de Jong, 1990). The expression construct includes a non-cleavable N-terminal His_6_ tag. The coding sequence for the recombinant protein is deposited in the SSGCID database (ID: TogoA.17199.a.B3.GE41306).

### Creating CvDCX1-mNeonGreenFP transgenic parasites in the TgDCX knockout background

The ptubg_CvDCX1-mNeonGreenFP plasmid was electroporated into *TgDCX* knockout parasites, which have been described previously (Nagayasu et al., 2016). The transfected population was subjected to five rounds of 20 µM chloramphenicol selection, and clones were individually screened for mNeonGreenFP fluorescence. The growth of the clones was compared with the *TgDCX* knockout, knock-in, and TgDCX complemented parasites using plaque assays.

### Light Microscopy

Wide-field image stacks were acquired using a DeltaVision imaging station (GE Healthcare / Applied Precision) fitted onto an Olympus IX-70 inverted microscope base as described in (Nagayasu et al., 2016). 3D-structured illumination (SIM) images were collected on a DeltaVision OMX Blaze system using a 100x NA 1.35 or 60X NA 1.42 objective with 3 orientations and 5 phases for the illumination pattern and a z-spacing of 0.125 µm. Image stacks were deconvolved with the manufacturer supplied software and locally measured wavelength-specific PSFs. *Contrast adjustment*: In the raw data, the range of reliably measured intensities often exceeds 200-fold. On a good monitor with linear mapping, perhaps a 30-fold range can be displayed. For print media, the range is even smaller. Therefore, with a linear mapping of data intensities to display media, it is not possible to display the biologically and statistically significant image data without making dim structures invisible or bright structures saturated. For this reason, we have routinely adjusted contrast levels with non-linear mapping (*i.e.,* gamma not equal to 1) to allow the reader to see both dim and bright features in cells of interest.

### Converting fluorescence intensity to number of molecules

Sindbis virus particles containing exactly 240 copies of a fluorescent protein were used as the calibration standard to convert the measured fluorescent intensity of cellular structures into the number of FP-tagged molecules in the structure. The preparation, extensive characterization and procedures for using these fluorescent virions for molecular counting are described in detail in (Murray, 2017). Briefly, images of virions adsorbed to a clean coverslip were recorded. After correcting for variations in illumination intensity across the field of view and subtracting background, the net fluorescence (photons/sec) due to single virus particles (240 fluorescent protein molecules) can be determined. Different preparations of virus have a variable content of dimers and typically a very small number of larger aggregates. The single and double particles are evident as two incompletely resolved peaks in the histogram of measured intensities. The two peaks can be resolved by fitting a two-component Gaussian distribution to the observed histogram. The numerical/image analysis program Igor Pro (Wavemetrics Inc, Oregon, USA) was used for this fitting. To improve the stability of the least-squares estimates of component means, outliers were removed from the data by trimming the highest and lowest 1% of the observed particle net intensities before fitting.

Using imaging conditions identical to those used for the virus sample, the net fluorescence intensity of the target cellular structure was similarly measured. In the case of microtubules, the length of the measured fluorescent segment was also determined, thus allowing the calculation of the number of FP-tagged molecules per unit length. A canonical 13-protofilament microtubule contains 1625 tubulin dimers per micron, so the number of FP-tagged molecules per tubulin dimer can then be calculated, assuming that the target structure contained a single microtubule. Alternatively, if one assumes that every tubulin dimer is bound to one FP-tagged microtubule binding protein, the number of microtubules in the target can be estimated.

### Measuring radius of curvature

3D stacks of images of cells transfected with plasmids driving expression of mCherryFP-tagged microtubule binding proteins were acquired. The *Xenopus* cell line used stably expresses EGFP_α-tubulin. Z-projection of 3-5 slices (0.3 µm/slice) were displayed, and clearly visible single or bundled microtubules were identified in the images. The microtubules were divided into segments having curvature of constant sign, and three points were manually marked at the beginning, end, and approximate middle of each segment. Circular arc profiles were fitted to the three 3-points using the Semper (Saxton, 1996) image processing package, yielding the radius of curvature, angular span, and circumferential length of the segment. Segments with no apparent curvature were typically less than 200 pixels (∼ 20 µm) in length. Thus, in practice, the maximum radius of curvature that could be estimated (i.e., curvature amounting to 1 pixel deviation from a straight line over a span of 200 pixels) is of the order of 500 µm. In the few cases where the three points lay exactly on a straight line, an infinite radius of curvature would be calculated. To avoid numerical complications, perfectly straight line segments were therefore arbitrarily assigned a radius of 1000 µm, and these segments were excluded from the statistical estimates (mean and standard deviation) of radius of curvature for the population.

### Electron microscopy of whole mount Toxoplasma

Electron microscopy of detergent-extracted whole mount *Toxoplasma* cytoskeleton was performed as previously described in (Nagayasu et al., 2016).

### Electron microscopy of intact and disassembled conoids

Isolated conoids were prepared by first treating a suspension of ∼ 10^8^ extracellular parasites with 1 ml of a calcium-saline (138 mM NaCl, 2.7 mM KCl, 20 mM K^+^HEPES pH 7.6, 5 mM CaCl_2_) plus 25 µM A23187 to cause extension of the conoid (Mondragon and Frixione, 1996). After 5 min, the parasites were lysed by washing with 1 ml of detergent-containing buffers of decreasing ionic strength (first, 5 mM CHAPS [Sigma #C5070], 1 mM Tris acetate, pH 7.5, 10 mM K_2_EGTA; second, the same except 5 mM K_2_EGTA; third, two times with the same solution without K_2_EGTA). After each wash, the parasites were collected by centrifugation at 4500 x g for 5 min. The lysed parasites were resuspended in 2 ml of the final wash buffer in a 15 ml plastic tube and sonicated on ice twice for 30 sec, with cooling for 1 minute between pulses. The suspension was centrifuged at 13,200 x *g* for 5 min and the supernatant was discarded. The barely visible pellet, which contained mostly isolated conoids plus intact or fragmented apical polar rings, was resuspended in 100 µl 20 mM K^+^HEPES, pH 7.5. To prepare dissociated conoid fibers, the conoid prep was treated with 10 µg/ml α-chymotrypsin at room temperature for 6 min. Digestion was stopped by addition of TPCK (Sigma # T4376) to 100 µM. For cryoEM, a few µl of suspension was placed on an EM grid coated with a lacey carbon film, blotted with filter paper, and immediately frozen by plunging into a rapidly-stirred liquid propane:isopentane mixture (∼19:1, v/v) held at liquid nitrogen temperature. Frozen grids were searched in a Phillips 400 EM for appropriate areas at low illumination and defocus of ∼0.8 mm. Low dose images were acquired at 28,000 magnification, at 120 kev, and defocus of 1.6 or 2.0 µm. For negative staining, the suspension was allowed to adsorb for 15 min to a carbon-coated EM grid, negatively stained with 2% phosphotungstic acid, and imaged in a JEOL 1010 at 80kev, or a JEOL1400 Plus at 120kev.

### Correlative Light and Electron microscopy of Xenopus S3 cells transfected with a plasmid expressing fluorescently tagged TgDCX or CvDCX1

3D stacks of LM images of suitable cells in a 35 mm glass-bottom dish were recorded as described above at 60x magnification. Lower magnification (20x, 10x, and 4x) images of the same area were then recorded to facilitate relocating the same cells after embedding for EM. The dish of cells was simultaneously partially fixed and permeabilized by replacing the culture medium with Buffer 1 (100 mM Na^+^MES, 2 mM MgCl_2_, 100 µM CaCl_2_, 1 mM CaEGTA, pH 6.6) plus 100 mM glutaraldehyde and 0.5% Triton X-100. After shaking for 1 minute at room temperature, the cells were further permeabilized with 0.5% Triton X-100 in Buffer 1 for 30 min, then fixed with 100 mM glutaraldehyde plus 1% tannic acid (Sigma # T0125) in Buffer 1 for 2-4 hrs. After carefully aspirating off all of the fixative, 40 mM OsO_4_ in 15 mM NaHPO_4_ pH 6.0 was added and the dish was shaken at 4 °C for 1 hour. The dish was rinsed three times with H_2_O, shaken with 1% uranyl acetate for 4 hr, then dehydrated for 5 min each with 50%, 70%, and 95% EtOH, then with 100% EtOH for 5 min twice. The sample was infiltrated with freshly prepared Spurr resin: acetone (1:3) for 1 hr, then 3:1 for 1 hour, and 100% Spurr resin for ∼16 hr. The resin was aspirated off, replaced with 400 µl of fresh resin, covered with a glass coverslip, and polymerized at 66 °C for ∼ 16 hr. Cells originally imaged by LM were relocated by bright-field microscopy at 10X magnification in this thin sheet of resin sandwiched between two glass coverslips, and their locations marked. The glass coverslips on both sides of the thin resin layer were dissolved using concentrated HF, preserving the marked locations, and the thin sheet of resin containing the monolayer of cells was cut into small pieces, each piece containing one of the originally imaged cells. These small pieces were then placed in a block-mold, carefully oriented to allow sectioning perpendicular to the plane of the cell monolayer, and with optimal rotational orientation of the target cell. After re-embedding each small piece in a block of Spurr resin, the block was carefully trimmed so that the mesa for sectioning contained only the desired cell with a small margin on each side. Silver sections were cut with a diamond knife, deposited on bare or carbon-coated Cu grids, stained for 30 min with 6% uranyl acetate in 50% methanol, then for 5-10 min with Reynold’s lead citrate, and examined in a JEOL1400 EM at 120kev.

### Expression of recombinant TgDCX

TgDCX expressed in bacteria, tagged with hexahistidine or fused to maltose-binding protein, could be solubilized only in the presence of denaturing detergents, GuCl, or urea. Attempted removal of the denaturing agents and refolding by a variety of methods always resulted in precipitation of the TgDCX. The same is true of a TgDCX fragment containing the partial P25α domain and the DCX domain (*i.e*., lacking the first 70 aa). Expression of His_6_-tagged TgDCX in yeast (*K. lactis*), or of HAT-tagged mNeonGreenFP-TgDCX in HEK293E cells driven by the CHEF1 promoter also gave no soluble protein under non-denaturing conditions. An initial small scale expression of His_6_-mNeonGreenFP-TgDCX in Sf9 cells yielded some protein with microtubule binding and bundling activities, but the purity was poor. Subsequent larger scale purifications yielded no soluble full-length TgDCX using physiological buffers for extraction. Abundant soluble recombinant TgDCX could be obtained by extracting the Sf9 cell pellet with 0.3 M CHAPS or Zwittergent 3-10, plus 3 M GuCl or stronger denaturing buffers, but removal of the detergent/denaturant inevitably led to immediate irreversible precipitation of all the TgDCX.

### Purification and Crystallization of recombinant His-tagged TgDCX148-243

His-tagged recombinant TgDCX148-243 (SSGCID ID: TogoA.17199.a.B3.PW38274) was expressed in *E. coli* Rosetta BL21 (DE3) transformed with the plasmid BG1861_ TgDCX148-243 following standard SSGCID protocols as described previously (Choi et al., 2011). Purification was completed using Ni-NTA affinity and size exclusion chromatography following standard SSGCID protocols (Bryan et al., 2011). The purified protein was concentrated to 15 mg/mL in its final buffer (25 mM HEPES pH 7.0, 500 mM NaCl, 5% glycerol, 2 mM DTT, and 0.025% sodium azide), flash frozen in liquid nitrogen, and stored at −80°C. All crystallization experiments were completed in 96-well XJR trays (Rigaku Reagents) with 0.4 µl protein and 0.4 µl reservoir solution as sitting drops equilibrated against 80 µL reservoir. Crystallization trays were incubated at 14°C. Diffraction quality crystals were obtained in PACT Premier HT-96 (Molecular Dimensions) condition E6 (18% w/v PEG3350, 200 mM sodium formate). Experimental phases were obtained by soaking these crystals in reservoir containing 20% of a 2.5 M sodium iodide solution in ethylene glycol as cryoprotectant before vitrification by dipping in liquid nitrogen. The phasing information from this crystal was then applied to data collected from a separate crystal obtained using an optimization screen based on PACT E6 (24.09% w/v PEG3350, 50 mM sodium formate), which was soaked in 20% ethylene glycol as cryoprotectant and vitrified by dipping in liquid nitrogen.

### Data Collection and Structure Solution of recombinant TgDCX148-243

Data sets were collected in-house at 100 K on a Rigaku FR-E^+^ SuperBright rotating anode equipped with Rigaku VariMax optics and a Saturn 944+ CCD detector, using CuKα X-rays. All data sets were reduced with the XDS (Kabsch, 2010) (Supplementary Table 1). For the iodide data set, Friedel pairs were kept separate. Friedel pairs were merged for the data set used for refinement. For the iodide-soaked crystal, 11 anomalous sites were found with HySS (Grosse-Kunstleve and Adams, 2003) using data up to 2.00 Å resolution. The anomalous sites were further refined, and initial phases were calculated with Phaser_EP (Read and McCoy, 2011) within the CCP4 package (Collaborative Computational Project, 1994). The CCP4 program PARROT (Cowtan, 2010) was used for phase improvement; NCS averaging was not used due to low NCS correlations. An initial model was built with ARPwARP (Langer et al., 2008).

### Structure Refinement and Validation of recombinant TgDCX148-243

Molecular Replacement using the experimentally-phased model was completed with Phaser (McCoy et al., 2007) within the Phenix ligand pipeline. All structures were refined in PHENIX (Adams et al., 2010). Manual model building was completed using Coot (Emsley et al., 2010). The quality of all structures was assessed using built-in tools in Coot and using Molprobity (Chen et al., 2010) through the Phenix interface. The coordinates and structure factors were deposited in the PDB under accession code (<u>6B4A</u>).

### Structural modeling of TgDCX orthologues based on the TgDCX148-243 crystal structure

Homology models for TaDCX, PfDCX, VbDCX1, VbDCX3, CvDCX1, and CvDCX3 were obtained using Robetta (<u>robetta.org</u>) with the TgDCX structure as the top template and comparative models generated using the RosettaCM protocol (Song et al., 2013). As the sequence homology to their respective paralog is greater than 80%, CvDCX2 and VbDCX2 were threaded with Modeller against the Robetta coordinates for CvDCX3 and VbDCX3, respectively, and the models thus obtained were energy minimized against the Amber force field with 100 steps of steepest descent followed by 10 steps of conjugate gradient minimization (Webb and Sali). Electrostatic potential maps were calculated with APBS and displayed on the molecular surface with UCSF Chimera, with the ±10 kT/e potential isocontours shown as blue and red (Baker et al., 2001; Pettersen et al., 2004). Prior to electrostatic potential calculations, the human DCX-N structure (PDB:<u>5IOI</u>) was altered in UCSF Chimera by replacing the mutated residues D134-D135 with native lysines using the Dunbrack rotamers library (Burger et al., 2016; Dunbrack, 2002). The orientation relative to tubulin of the TgDCX and human DCX-C (PDB:<u>5IP4</u>) structures and homology models was estimated by superimposing the coordinates on the human DCX-N structure in complex with tubulin (PDB: <u>4ATU</u>) (Liu et al., 2012).

## Availability of data and materials

All data generated or analysed during this study are included in this published article and its additional files.

## ACKNOWLEDGEMENTS

We thank Drs. John Daum and Gary Gorbsky (Oklahoma Medical Research Foundation, Oklahoma City, OK) for the EGFP-tubulin *Xenopus* S3 cell line, Dr. Richard Day (Indiana University School of Medicine, Indianapolis, IN) for the pmNeonGreen-N1 plasmid, Dr. David Walliker for *Plasmodium falciparum*-3D7A genomic DNA (obtained through BEI Resources, NIAID, NIH), Yulia Pivovarova (Max F. Perutz Laboratories, Vienna, Austria) and Dr. Christopher Nosala (Indiana University) for insightful discussions, Dr. Amanda Rollins and Qing Zhang for technical support. We thank Christiane Hassel of the Indiana University Bloomington (IUB) Flow Cytometry Core Facility for assistance with flow cytometry, Dr. James Powers of the IUB Light Microscopy Imaging Facility for assistance and support with light microscopy (NIH S10-RR028697), and Drs. Barry Stein and Che-Yen Wang of the IUB Electron Microscopy Center for technical support and advice with electron microscopy. We are indebted to Dr. Owen Saxton (University of Cambridge) for providing the source code for his ‘Semper’ image processing package. This study was supported by funding from the American Heart Association (Postdoctoral Fellowship, 18POST34090005) awarded to JML, NIH-NIAID (R01-AI132463) and the March of Dimes (6-FY18-674) awarded to KH, and facility funding from the Indiana Clinical and Translational Sciences Institute to KH, funded in part by grants #UL1 TR001108 and #TL1TR001107 from the National Institutes of Health, National Center for Advancing Translational Sciences, Clinical and Translational Sciences Award. This project has also been funded in part with Federal funds from the NIH-NIAID under Contract No.: HHSN272201700059C.

---

**Supplementary Figure 1.**
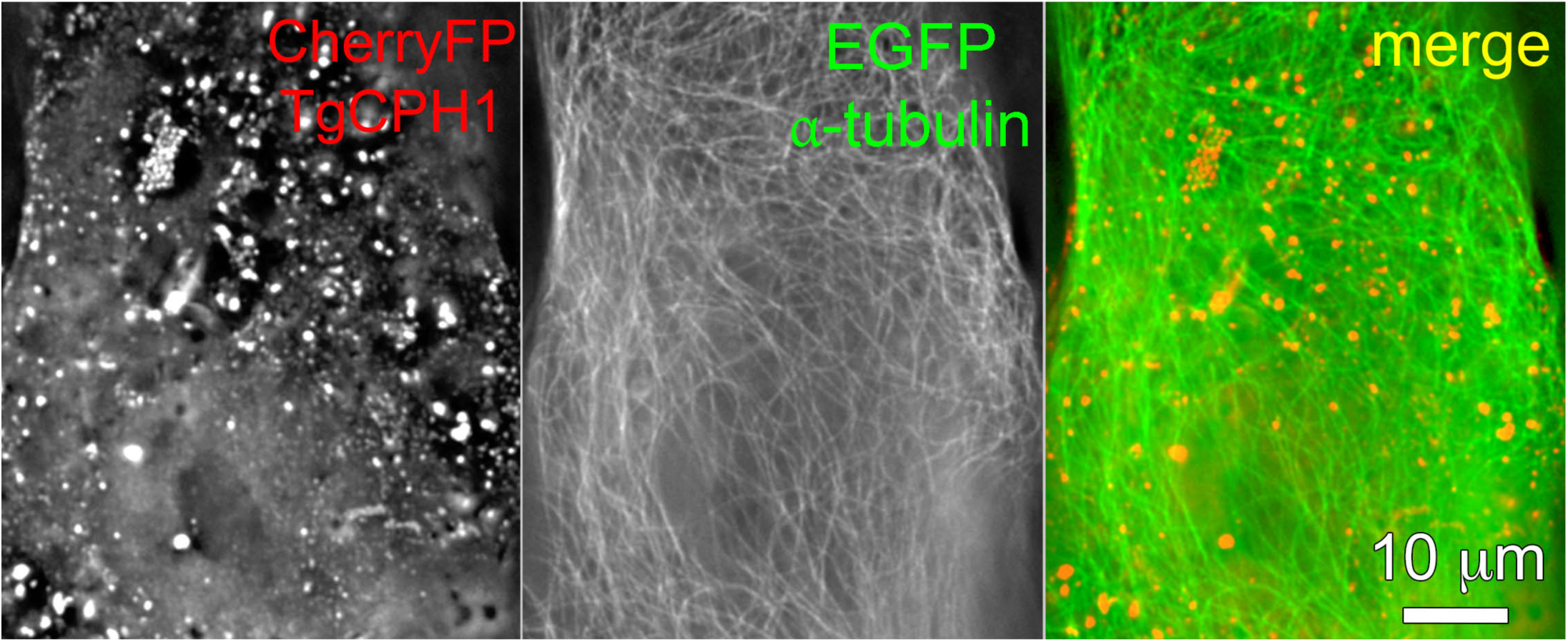
TgCPH1 does not bind to microtubules in the absence of TgDCX in *Xenopus* S3 cells. Deconvolved wide-field images of *Xenopus* S3 cells expressing mCherryFP-TgCPH1 (*red*), and EGFP-α-tubulin (*green*). When expressed alone, FP tagged CPH1 is diffusible cytoplasmic and in vesicles, not associated with microtubules.

**Supplementary Figure 2.**
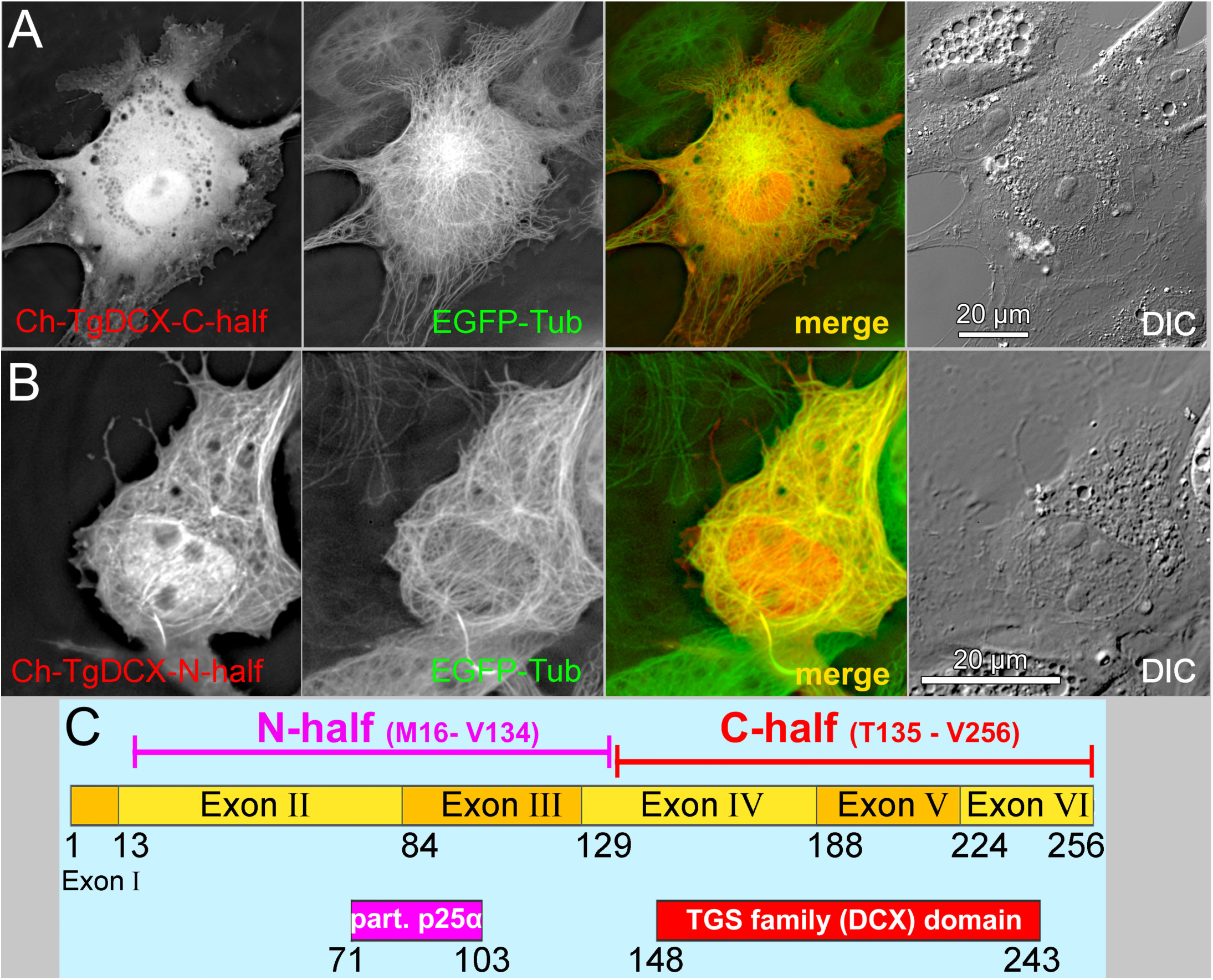
Effect of single domain constructs on microtubules in *Xenopus* S3 cells. A) TgDCX135-256, containing the C-terminal half of TgDCX, which includes the DCX domain, remains cytoplasmic, not binding to microtubules. B) TgDCX16-134, containing the N-terminal half of TgDCX, which includes the partial P25α domain, binds to microtubules in *Xenopus* S3 cells but does not modify MT architecture. C) Schematic showing the N-half and C-half fragments of TgDCX.

**Supplementary Figure 3.**
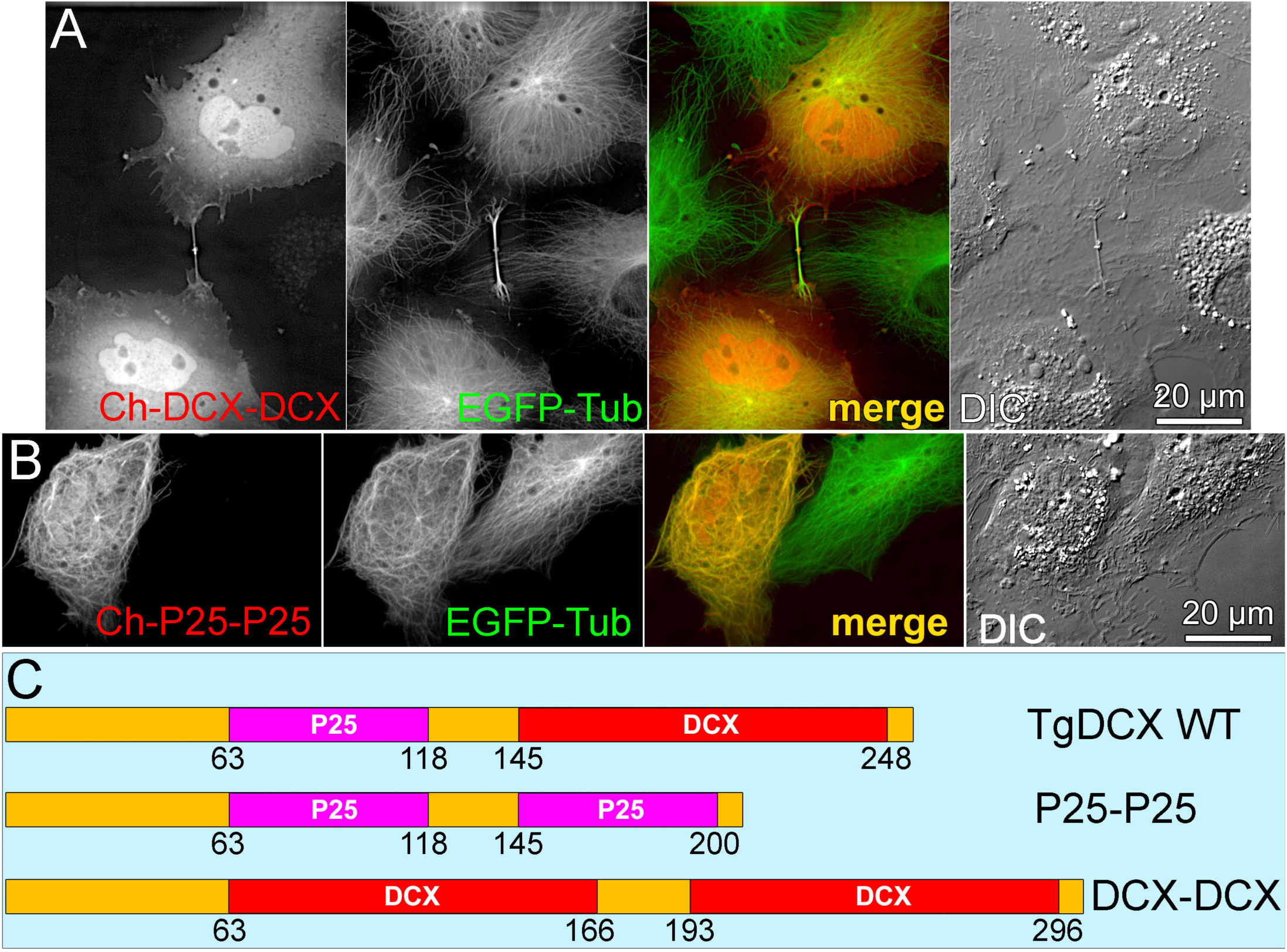
Effect of domain swapping on TgDCX interaction with microtubules in *Xenopus* S3 cells. A) A construct with two copies of the DCX domain remains cytoplasmic, not binding to microtubules. B) A construct with two copies of the partial P25α domain binds to microtubules but does not change their architecture. C) Domain structure of the WT TgDCX and the domain-swapped constructs. The sequences outside the defined domains, shown as orange segments, are identical in all three molecules.

**Supplementary Figure 4.**
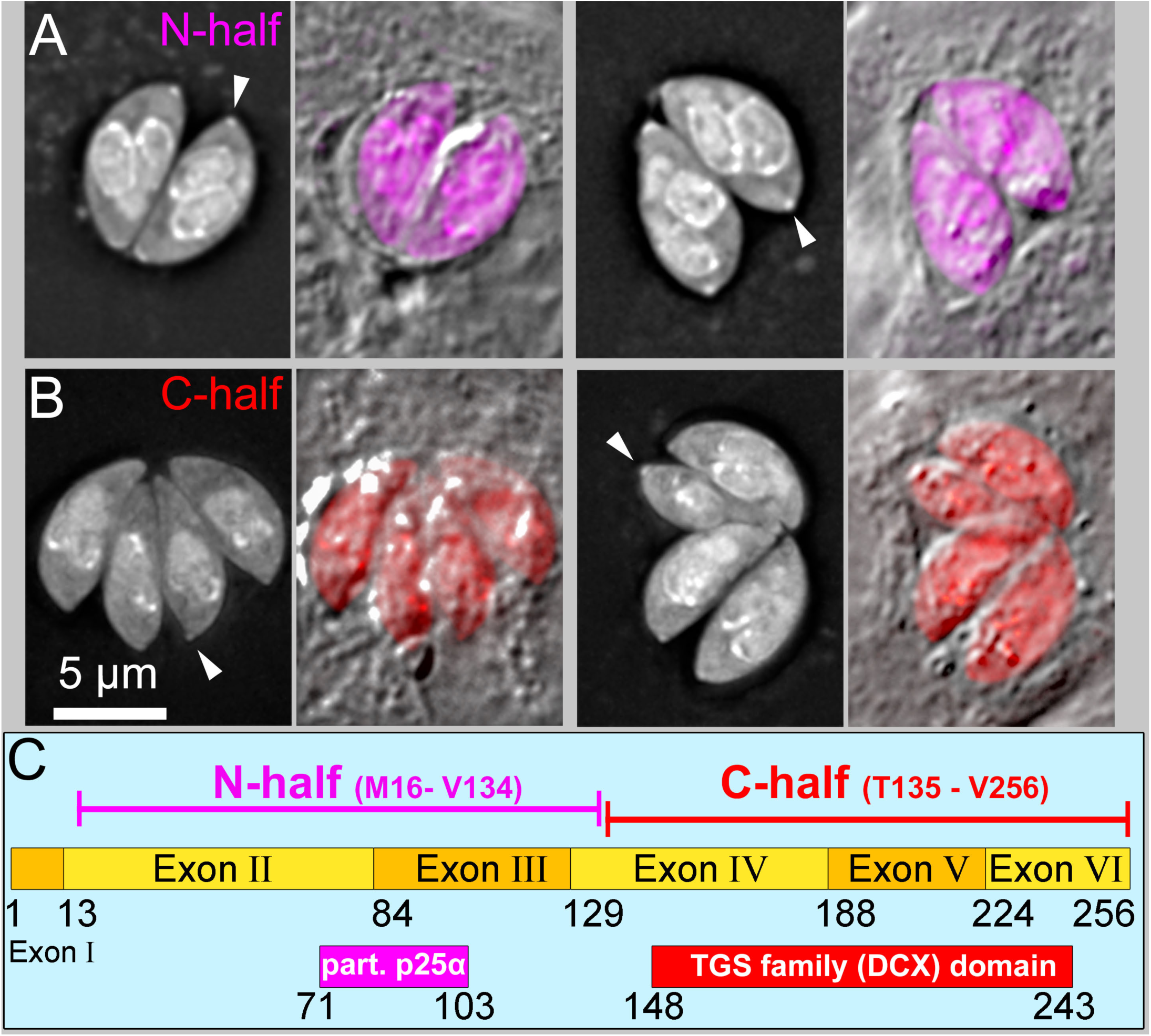
Deconvolved wide-field images of parasites expressing either eGFP-TgDCX16-134 (“N-half”, A) or eGFP-TgDCX135-256 (“C-half”, B). Arrowheads indicate conoids. (C) Diagram of the domain structure of TgDCX. Numbers indicate amino acid residues. The full-length protein is 256 aa.

**Supplementary Figure 5.**
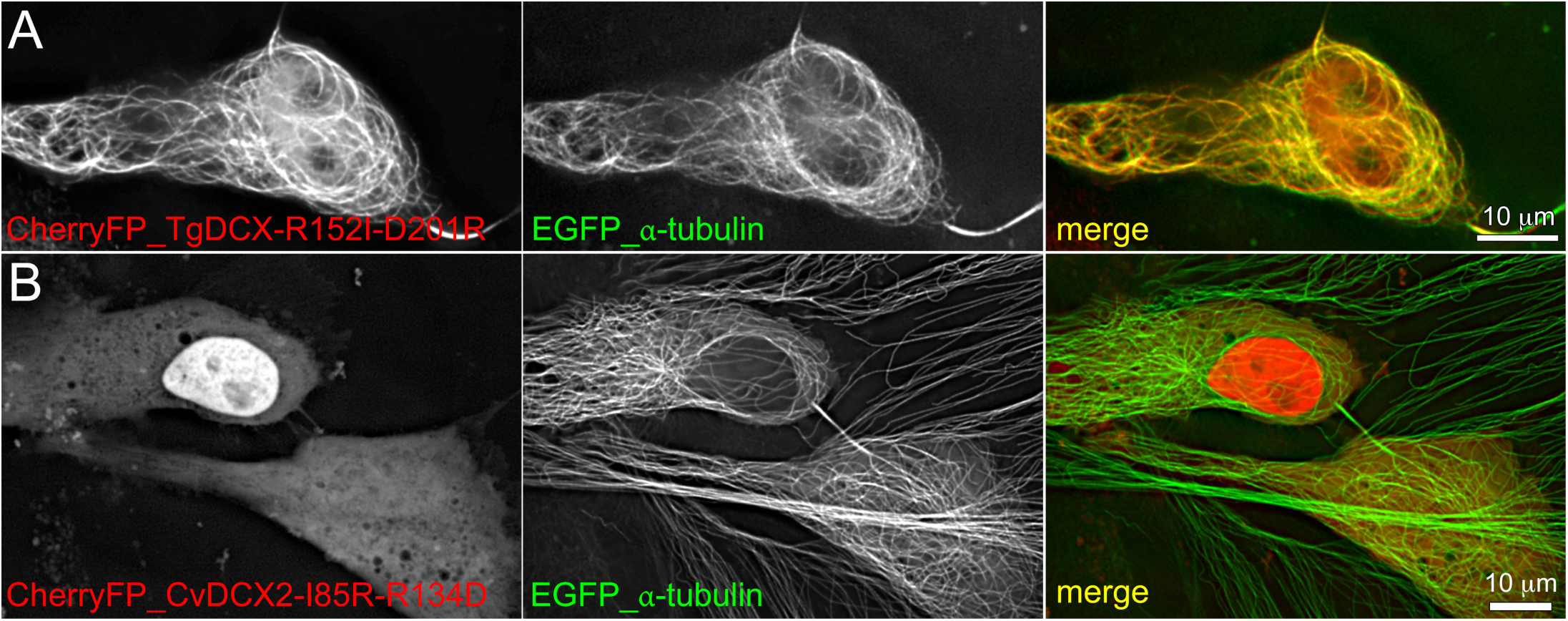
Residues R152 and D201 in TgDCX do not affect microtubule binding or curving in *Xenopus* S3 cells. Deconvolved wide-field images of *Xenopus* S3 cells expressing EGFP-tubulin (*green*) and either mCherryFP-TgDCX-R152I_D201R (A, *red*) or mCherryFP-CvDCX2-I85R_R134D (B, *red*).

**Supplementary Table 1:** Data Collection, Phasing, Ramachandran Statistics, and Refinement Statistics for TogoA.17199.a.B3.PW38274

| Parameters | Iodide | Native |
| --- | --- | --- |
| Wavelength (Å) | 1.5418 | 1.5418 |
| Space Group | P2<sub>1</sub>2<sub>1</sub>2<sub>1</sub> | P2<sub>1</sub>2<sub>1</sub>2<sub>1</sub> |
| *Cell Dimensions* |  |  |
| *a*, *b*, *c* (Å) | 36.40, 53.28, 96.25 | 36.86, 53.23, 96.38 |
| $\alpha$, $\beta$, $\gamma$ (°) | 90.000, 90.000, 90.000 | 90.000, 90.000, 90.000 |
| Resolution (Å) | 50.00-2.00 | 50.00-2.00 |
| *R*<sub>merge</sub> (%) | 4.5 (14.4) | 5.9 (55.7) |
| *I*/σ*I* | 44.1 (14.2) | 18.0 (2.7) |
| Completeness (%) | 98.7 (88.5) | 99.8 (99.8) |
| Redundancy | 14.0 (8.5) | 6.1 (3.9) |
| *Refinement* |  |  |
| Resolution (Å) |  | 35.7-2.0 |
| Reflections (#) |  | 13372 |
| *R*<sub>work</sub>/*R*<sub>free</sub> (%) |  | 18.6/26.1 |
| MolProbity Score |  | 1.3 |
| Ramachandran Favored |  | 97.8% |
| Ramachandran Allowed |  | 2.2% |
| Ramachandran Outliers |  | 0.0% |
| Number of Non-Hydrogen Atoms |  |  |
| Protein |  | 1463 |
| Formate |  | 6 |
| Water |  | 104 |
| *R*<sub>msds</sub> |  |  |
| Bond Lengths (Å) |  | 0.007 |
| Bond Angle (°) |  | 0.784 |
| Average B Factors (Å<sup>2</sup>) |  |  |
| Overall |  | 45.67 |
| Protein |  | 45.59 |
| Formate |  | 55.3 |
| Water |  | 45.29 |

**Supplementary Table 2.**
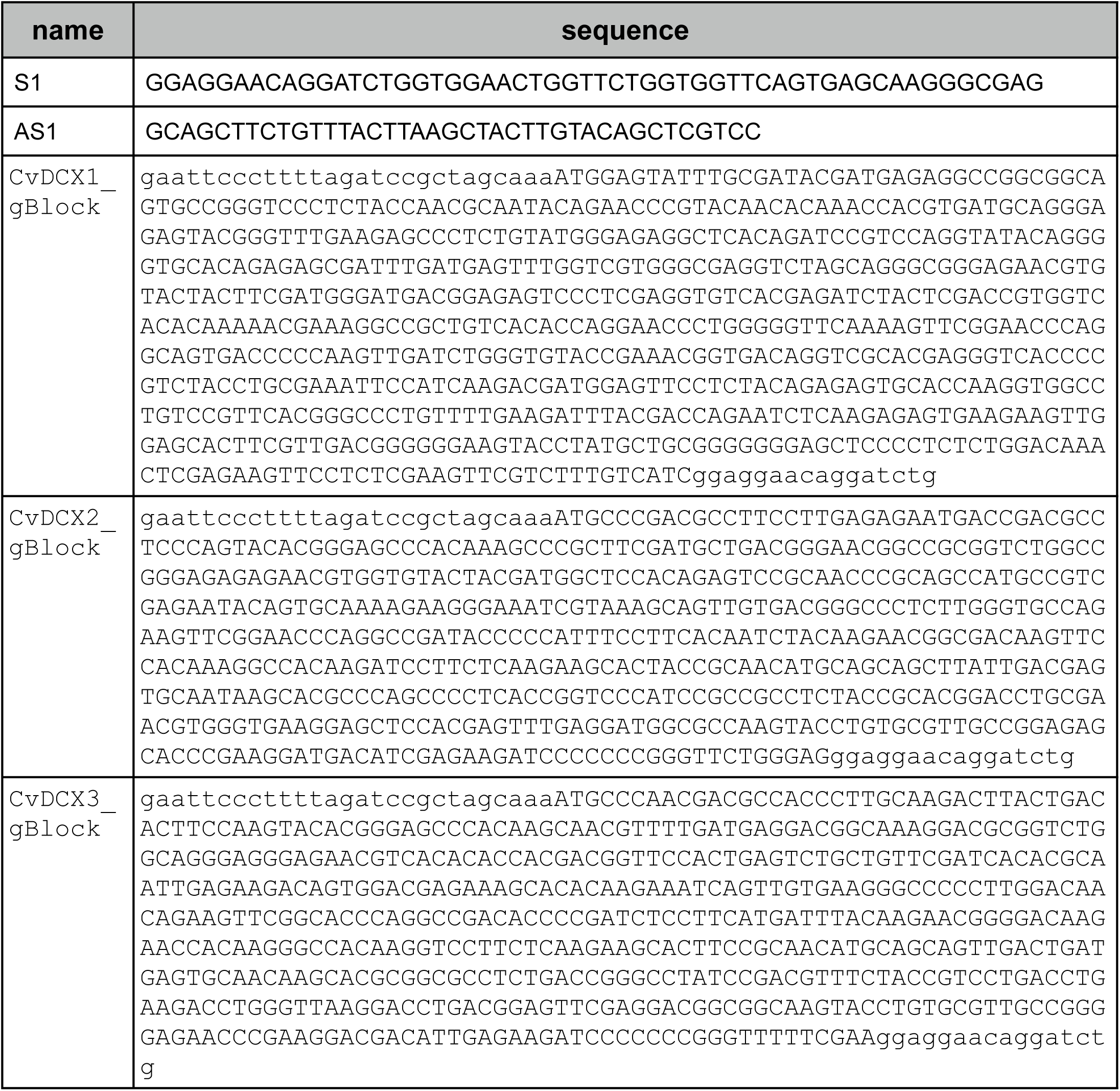
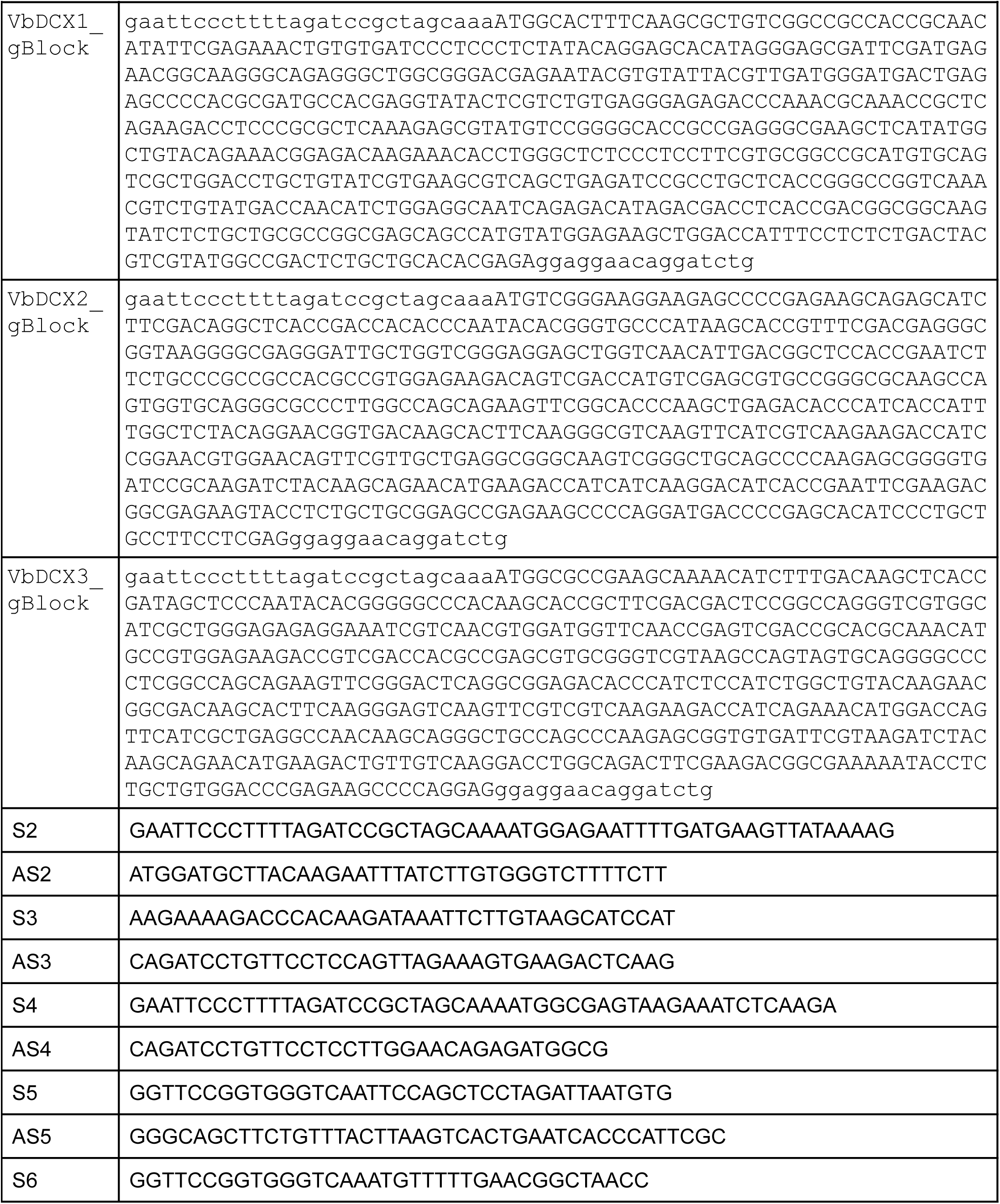
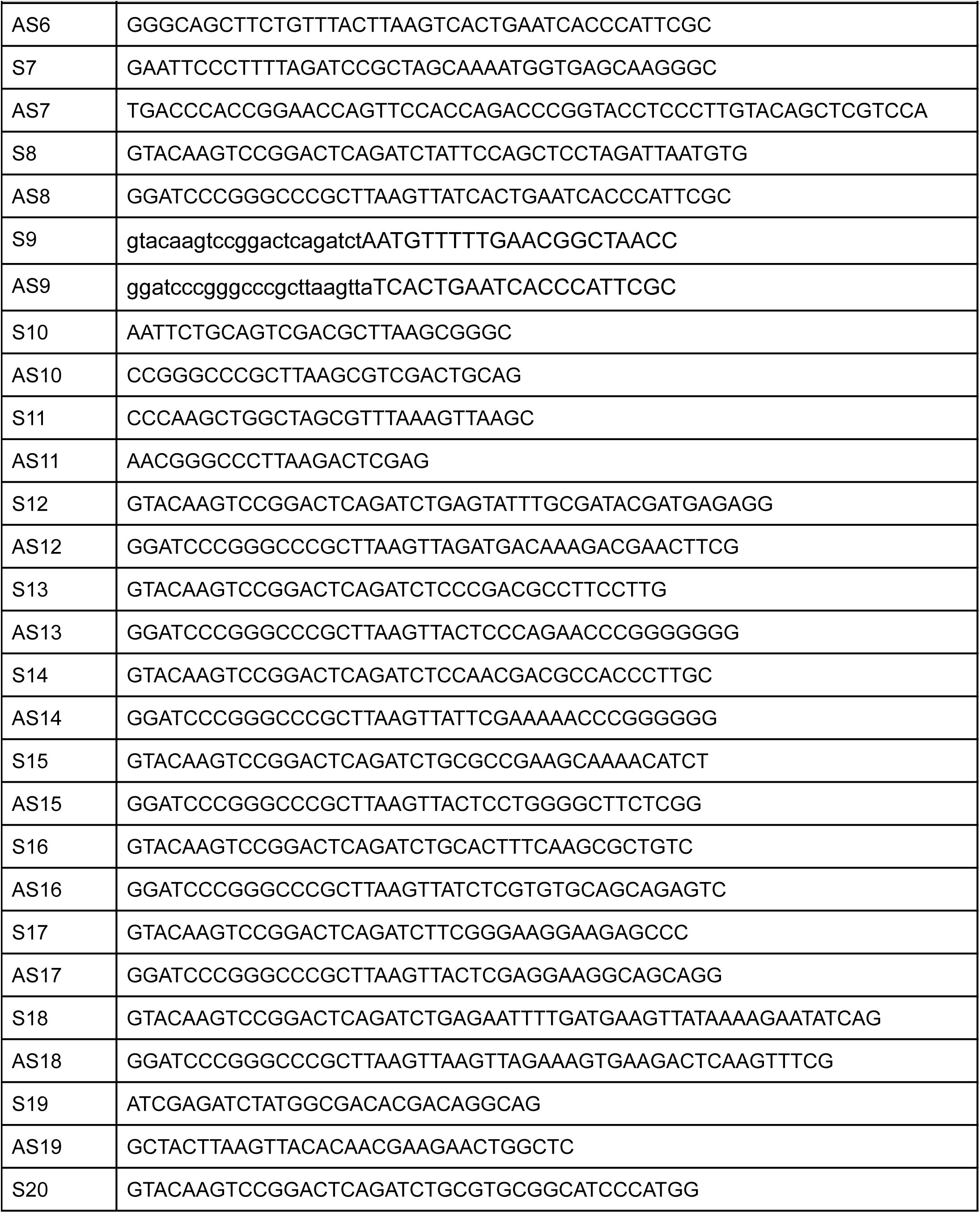
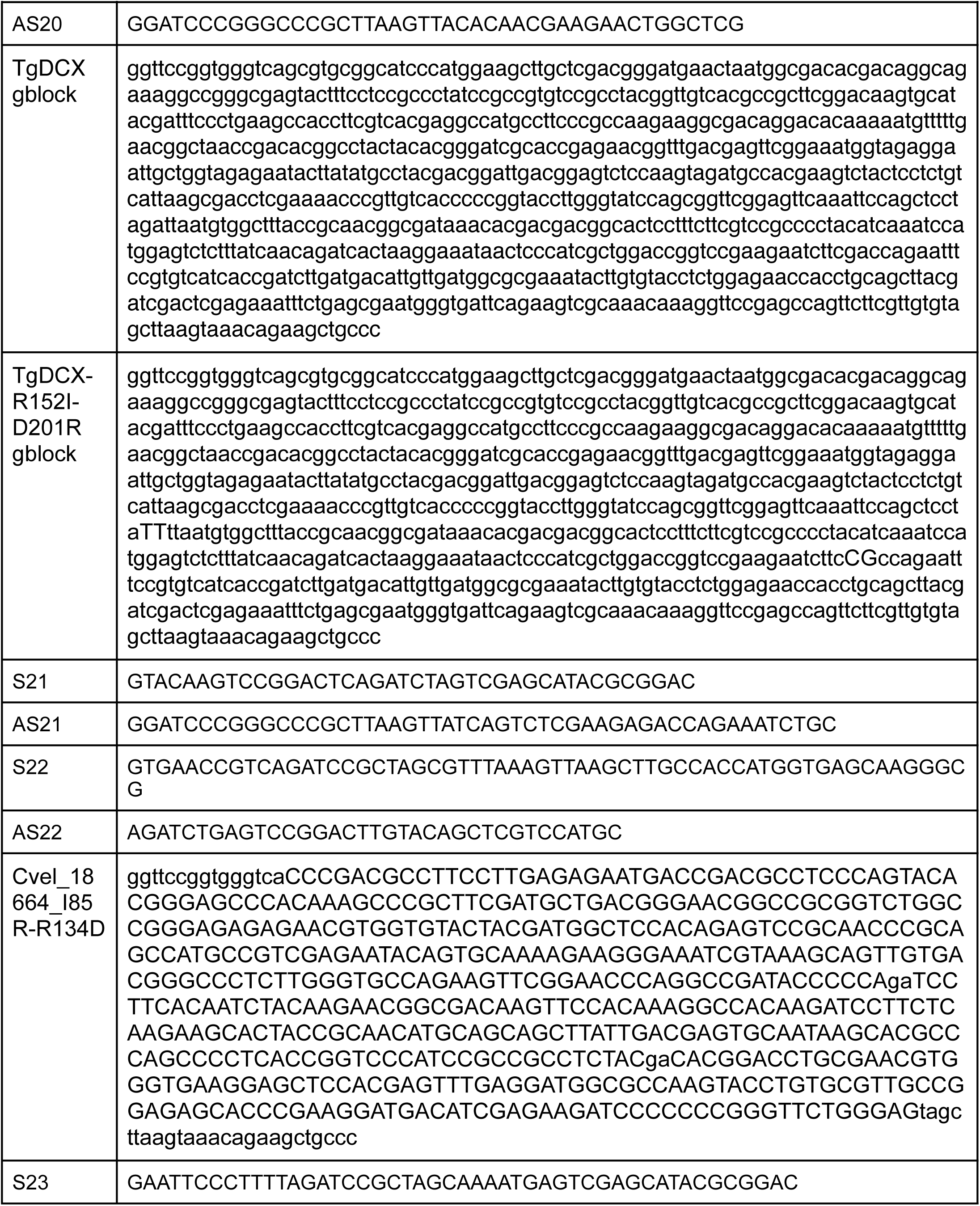
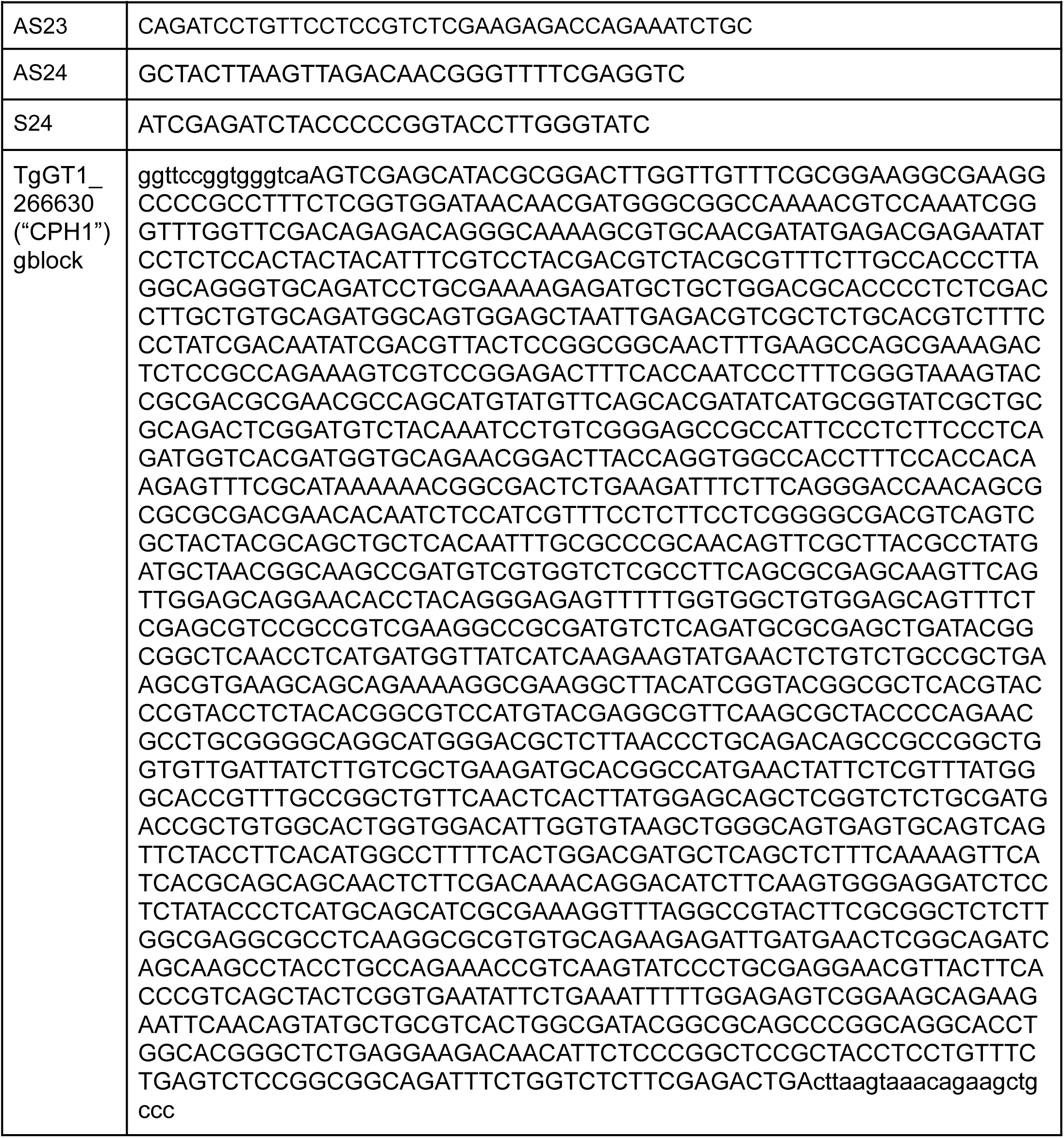
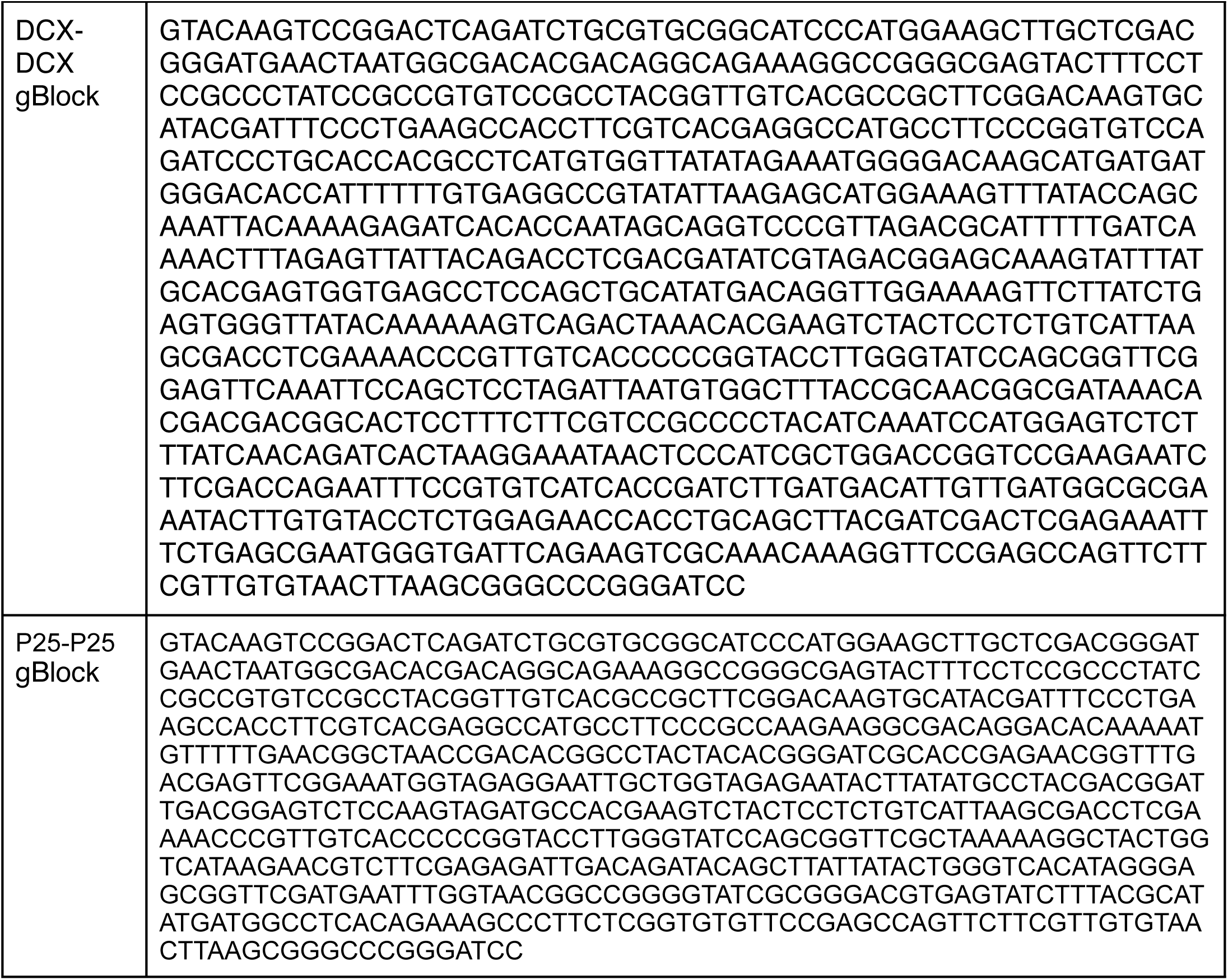
Oligonucleotides and gBlock fragments used in this study

**Supplementary movie 1.** A 3-D stack of deconvolved wide-field images of *Xenopus* S3 cells expressing EGFP-α-tubulin (*green*) and TgDCX-mCherryFP (*red*). Bar: 10 μm.

**Supplementary movie 2.** A 3-D stack of deconvolved wide-field images of *Xenopus* S3 cells expressing EGFP-α-tubulin (*green*) and mCherryFP-TgDCX (*red*). Bar: 10 μm.

**Supplementary movie 3.** A 3-D stack of deconvolved wide-field images of *Xenopus* S3 cells expressing EGFP-tubulin (*green*) and mCherryFP-TgDCX71-243 (*red*). Bar: 10 μm.

